# The malaria parasite sheddase SUB2 governs host red blood cell membrane sealing at invasion

**DOI:** 10.1101/2020.07.15.205732

**Authors:** Christine R Collins, Fiona Hackett, Steven A Howell, Ambrosius P Snijders, Matthew RG Russell, Lucy M Collinson, Michael J Blackman

## Abstract

Red blood cell (RBC) invasion by malaria merozoites involves formation of a parasitophorous vacuole into which the parasite moves. The vacuole membrane seals and pinches off behind the parasite through an unknown mechanism, enclosing the parasite within the RBC. During invasion, several parasite surface proteins are shed by a membrane-bound protease called SUB2. Here we show that genetic depletion of SUB2 abolishes shedding of a range of parasite proteins, identifying previously unrecognized SUB2 substrates. Interaction of SUB2-null merozoites with RBCs leads to either abortive invasion with rapid RBC lysis, or successful entry but developmental arrest. Selective failure to shed the most abundant SUB2 substrate, MSP1, reduces intracellular replication, whilst conditional ablation of the substrate AMA1 produces host RBC lysis. We conclude that SUB2 activity is critical for host RBC membrane sealing following parasite internalisation and for correct functioning of merozoite surface proteins.

**Key highlights:**

- Many malaria parasite surface proteins are shed by SUB2 during RBC invasion
- SUB2-null merozoites either induce rapid host RBC lysis, or invade then die
- Merozoite surface protein shedding is crucial for host RBC membrane sealing

## Introduction

The phylum Apicomplexa comprises a diverse group of protozoan organisms, many of which are obligate intracellular parasites of clinical or veterinary importance. A feature of these parasites is their possession of invasive forms that actively penetrate host cells. In the asexual blood stages of infection by malaria parasites (*Plasmodium* species), merozoites invade red blood cells (RBCs), replicating intracellularly to produce merozoites that egress to invade fresh RBCs. Invasion is a rapid, multi-step process that includes binding, merozoite reorientation, discharge of secretory organelles called rhoptries and micronemes, formation of an electron-dense ‘tight junction’ (TJ) between the merozoite apical end and the RBC membrane, and actinomyosin-powered entry into the RBC through this structure, with concurrent formation of a membrane-bound parasitophorous vacuole (PV) within which the parasite develops ^1, 2, 3, 4^. Invasion ends with sealing of the RBC behind the intracellular parasite, concomitant with pinching off of the nascent PV membrane (PVM) in a membrane scission event such that the PVM is eventually non-contiguous with and internal to the RBC membrane. Invasion is typically followed by transformation of the host RBC into a shrunken ‘spiky’ state called echinocytosis, which resolves within minutes. The parasite rapidly transforms into an amoeboid ‘ring’ form. Whilst many advances have been made over recent decades in understanding invasion, particularly in the most lethal malaria species *Plasmodium falciparum* and *P. knowlesi*, the mechanisms underlying PV formation are obscure and nothing is known of the molecular mechanism(s) responsible for host cell membrane sealing.

Early electron microscopic (EM) studies showed that invading merozoites shed a ‘fuzzy coat’ of ∼20 nm-long bristle-like fibres ^1, 4^. These are likely composed predominantly of the major glycosyl phosphatidylinositol (GPI)-anchored surface protein MSP1, which is synthesised as an abundant, ∼200 kDa precursor at the plasma membrane of developing parasites ^5^. Minutes before egress, MSP1 is proteolytically cleaved into several fragments by a parasite protease called SUB1 ^6, 7^. These form a noncovalent complex on the merozoite surface ^8^ where, together with several partner proteins, it facilitates membrane rupture at egress ^9^. During invasion the bulk of the MSP1 complex is shed from the merozoite ^10, 11, 12, 13^ as a result of a single further cleavage catalysed by a second, membrane-bound protease called SUB2 ^14, 15, 16^. SUB2 also mediates shedding of two merozoite surface integral membrane proteins called PTRAMP and AMA1, which are released from micronemes at around egress ^17, 18, 19, 20^. PTRAMP acts as an RBC binding ligand ^19^ while AMA1 plays a central role in TJ formation (reviewed in ^21^). To access its substrates, SUB2 is also released from micronemes onto the merozoite surface, where it translocates to the posterior pole of the parasite just before or during invasion ^10, 14^. Despite evidence that SUB2 is essential ^22, 23, 24^ and that shedding of MSP1 and AMA1 is important and can aid evasion of invasion-inhibitory antibodies ^12, 25, 26, 27^, the molecular function of SUB2-mediated shedding is unknown.

Here we used genetic modification of *P. falciparum* SUB2 and two of its substrates to examine the essentiality and function of SUB2 in the erythrocytic lifecycle. We show that SUB2 depletion results in defects in merozoite surface protein shedding and sealing of the host RBC upon invasion, leading to either abortive invasion with loss of host RBC haemoglobin, or developmental arrest of the intracellular parasite. This lethal phenotype results wholly or in part from a combination of lack of MSP1 shedding and loss of AMA1 function, highlighting SUB2 as a key mediator of parasite viability and host RBC membrane integrity.

## Results

### Efficient conditional ablation of SUB2 expression

The *P. falciparum sub2* gene encodes a type I integral membrane protein with a large ectodomain incorporating a subtilisin-like protease module ^15, 16^. We designed a construct to integrate into the *sub2* locus by homologous recombination, producing a modified locus in which the entire second exon encoding the crucial catalytic Ser961, transmembrane domain (TMD) and cytoplasmic domain was flanked (floxed) by *loxP* sites (Fig. 1A). Integration also fused a triple hemagglutinin (HA3) epitope tag to the SUB2 C-terminus, as achieved previously ^10, 14^. Transfection of the construct into the DiCre-expressing *P. falciparum* 1G5DC clone ^28^ resulted in outgrowth of WR99210-resistant parasites expressing HA3-tagged SUB2. Immunofluorescence analysis (IFA) of two integrant parasite clones (1B4int and FE5int) with anti-HA3 antibodies showed a signal consistent with the previously-determined location of SUB2 in micronemes, confirming correct gene modification (Fig. 1B).

**Fig. 1.**
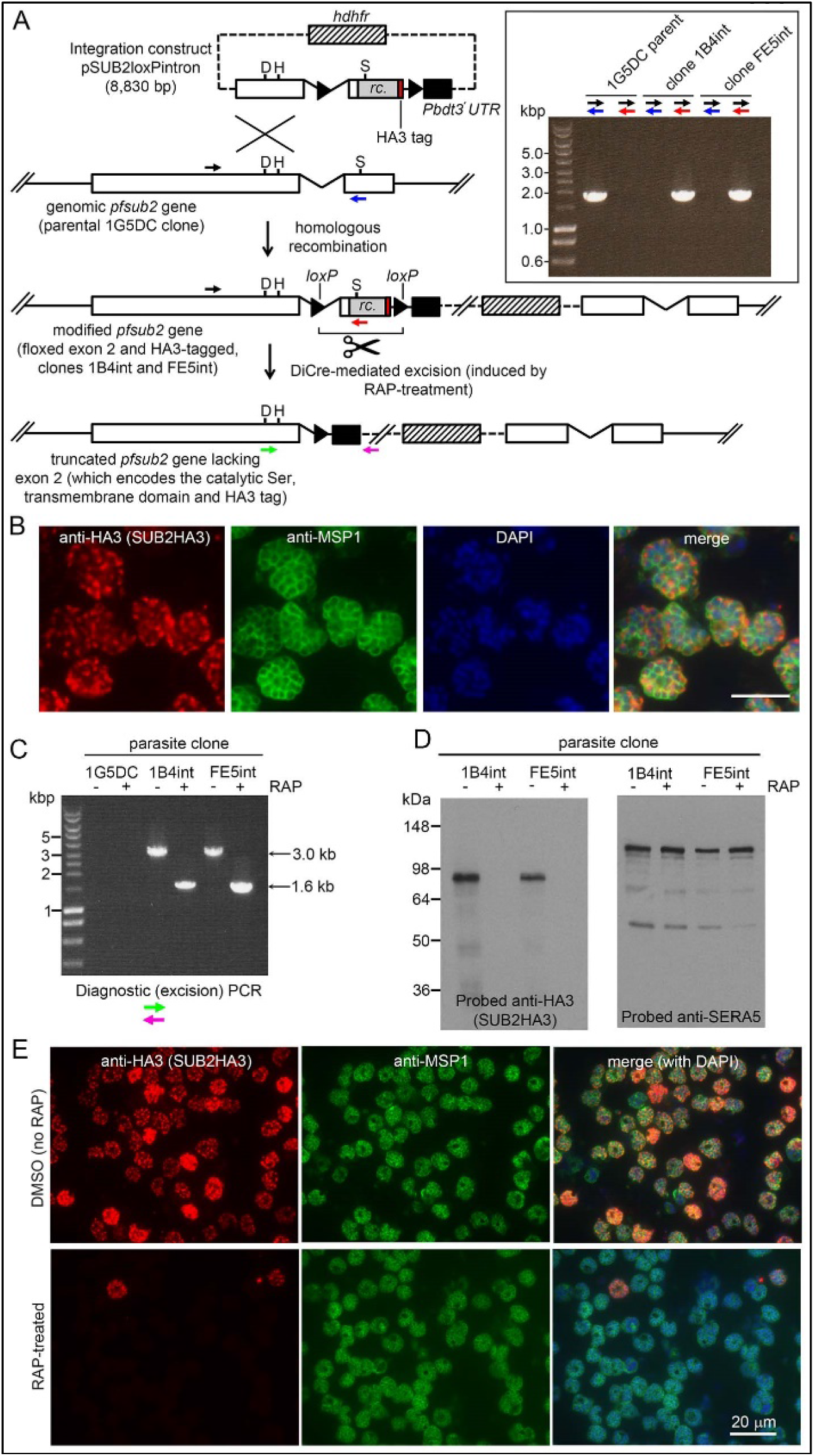
Conditional disruption of the *sub2* gene within a single erythrocytic cycle. (A) Floxing and epitope tagging. The *P. falciparum sub2* gene (PlasmoDB PF3D7_1136900) comprises two exons and a 143 bp intron. Targeting construct pSUB2loxPintron contains an internal ∼1,750 bp gene segment with the intron modified by insertion of a *loxP* site (arrow-head). This was fused to synthetic *sub2*_synth_ sequence ^14^ (rc.) encoding the TMD and cytoplasmic domain, plus an HA3 tag and stop codon, followed by a second *loxP* site then the 3’ UTR of the *P. berghei* dihydrofolate reductase (*dhfr*) gene (Pbdt3’ UTR, black box) to ensure correct gene transcription and polyadenylation. The human *dhfr* cassette confers resistance to the antifolate WR99210. Plasmid backbone, dotted line. Single-crossover recombination introduces the entire construct into the genome, floxing exon 2 and producing a promoterless downstream partial gene duplication. Catalytic triad residues; D, H, S. Colored arrows; primers for diagnostic PCR (see Methods for sequences and colour codes). Inset, diagnostic PCR confirming pSUB2loxPintron integration. Predicted sizes of PCR amplicons: 1,943 bp (black/blue primers); and 1,954 bp (black/red). (B) IFA localising SUB2HA3 in 1B4int schizonts. Parasite nuclei were stained with 4,6-diamidino-2-phenylindole (DAPI, blue). Scale bar, 10 μm. (C) PCR confirming RAP-induced excision of the floxed exon 2. (D) Western blot of schizont extracts showing loss of SUB2HA3 following RAP treatment. The PV protein SERA5 acted as loading control. (E) IFA showing loss of SUB2HA3 in cycle 0 1B4int schizonts following RAP-treatment of rings. Expression was detectable in just 2 ±0.1% of RAP-treated schizonts (the microscopic field was deliberately chosen to include 2 of the rare HA3-positive schizonts). See also Supplementary Fig. 1.

Excision of the floxed sequence was predicted to produce a truncated gene product lacking a functional catalytic domain, TMD and HA3 tag. To assess the efficiency of DiCre-mediated gene disruption, synchronised rings of both integrant clones were pulse-treated with rapamycin (RAP) which induces Cre recombinase activity. The parasites were examined ∼44 h later at the multinucleated schizont stage (when SUB2 expression is maximal) within the erythrocytic cycle of RAP-treatment (cycle 0). Diagnostic PCR showed efficient excision of the floxed *sub2* sequence (Fig. 1C), whilst Western blot confirmed virtually complete loss of the HA3 signal (Fig. 1D) and no signal was detectable by IFA in most of the RAP-treated schizonts (Fig. 1E). No defect in merozoite biogenesis was evident (Supplementary Fig. 1). These results confirmed disruption of SUB2 expression within a single erythrocytic cycle, and suggested that loss of SUB2 during intracellular development had no impact on schizont maturation.

### SUB2 is the merozoite surface sheddase; loss of SUB2 has no effect on egress but reduces merozoite surface protein shedding and RBC invasion

SUB2 is discharged onto the free merozoite surface to cleave its substrates ^10, 14, 17, 18^. We expected egress to be unaffected by loss of SUB2, and in accord with this time-lapse video microscopy showed no differences in rupture of mock-and RAP-treated 1B4int and FE5int cycle 0 schizonts (Fig. 2A). To assess the invasive capacity of the released merozoites, highly synchronised schizonts were incubated for 4 h with RBCs to allow egress and invasion. Newly-invaded (cycle 1) rings were produced in the ΔSUB2 cultures, but at levels only ∼50% of those in control cultures (Fig. 2B). Western blot analysis of medium harvested from the ΔSUB2 cultures showed substantially decreased levels of MSP1, AMA1 and PTRAMP, consistent with loss of shedding (Fig. 2C). SUB2-mediated shedding during invasion is highly efficient, so antibodies specific to shed segments of MSP1 and AMA1 are invariably unreactive with newly-invaded rings ^11, 12, 13, 29^. In contrast, the membrane-proximal AMA1 ‘stub’ and GPI-anchored MSP1_19_ domain that remain on the parasite surface following cleavage at the respective juxtamembrane sites are readily detected in rings ^12, 13, 17, 25^. We therefore used selected antibodies to examine the cycle 1 rings by IFA. Monoclonal antibodies (mAbs) X509 and 89.1, which recognize shed fragments of the MSP1 complex, did not react with control rings, as expected. In contrast, both mAbs strongly recognised the ΔSUB2 cycle 1 rings (Fig. 2D), indicating that in the absence of SUB2, unshed MSP1 complex was carried into RBCs on invading merozoites. Similarly, antibodies to the AMA1 ectodomain showed stronger reactivity with ΔSUB2 rings than with control rings (Fig. 2E).

**Fig. 2.**
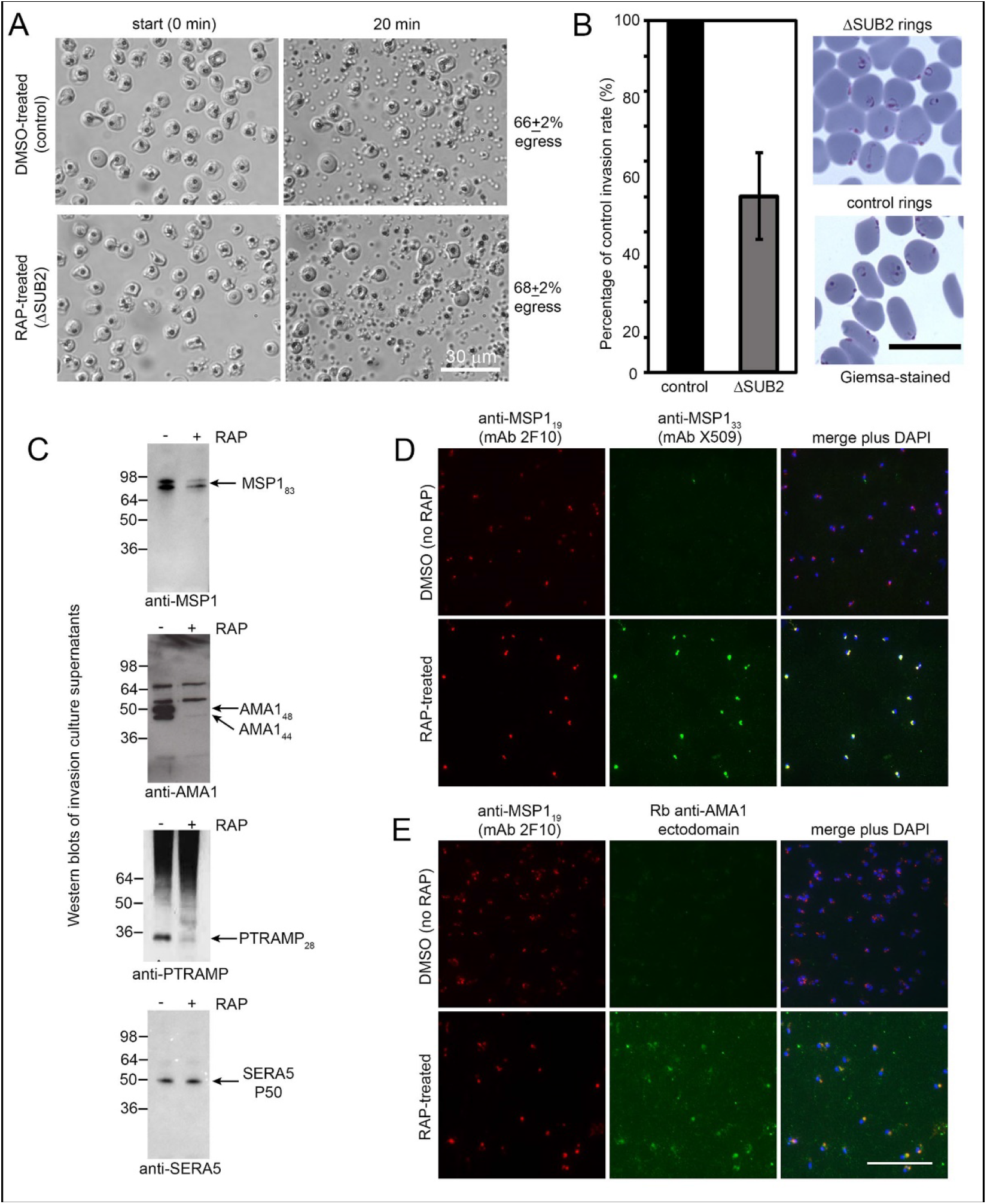
SUB2 is not required for egress, but is implicated in invasion and is required for shedding of MSP1, AMA1 and PTRAMP. (A) DIC microscopic images showing rupture of control and ΔSUB2 FE5int schizonts. (B) Invasion assay showing ∼50% reduction of invasion in ΔSUB2 1B4int parasites. Data are presented as percentage of ring formation by control parasites (mean +/− SD of 3 biological replicate experiments). Ring parasitaemia in control cultures ranged from 11.5-45.5%. Right, Giemsa-stained cycle 1 rings, 4 h following invasion. (C) Western blot of culture supernatants following incubation of control and ΔSUB2 1B4int schizonts with fresh RBCs for 4 h. Shed proteolytic fragments are indicated. Release of SERA5, which is not processed by SUB2, acts as a loading control. (D) IFA of rings produced following invasion by control and ΔSUB2 parasites. Only in the ΔSUB2 rings are the MSP1 and AMA1 ectodomain detectable. All rings are reactive with the anti-MSP1_19_ mAb 2F10. Scale bars, 20 μm

To interrogate the global effects of SUB2 disruption, culture media harvested following rupture of control and ΔSUB2 schizonts were analysed by quantitative mass spectrometry. This revealed reduced levels of several merozoite surface proteins in the ΔSUB2 egress supernatants, consistent with their reduced shedding (Fig. 3 and Supplementary Table 1). The depleted proteins included SUB2 itself, as well as AMA1, PTRAMP and members of the MSP1 complex (MSP1, 3, 6 and 7), all in accord with the Western blot and IFA data. Unexpectedly, the most depleted proteins also included the GPI-anchored merozoite surface proteins Pf92, MSP2, MSP4 and MSP5 ^30^ as well as the MSP7-like protein MSRP2, none of which are components of the MSP1 complex ^31, 32, 33, 34, 35^. These results suggest that SUB2 is required to shed these proteins too, indicating a broader repertoire of merozoite surface substrates than previously appreciated.

**Fig. 3.**
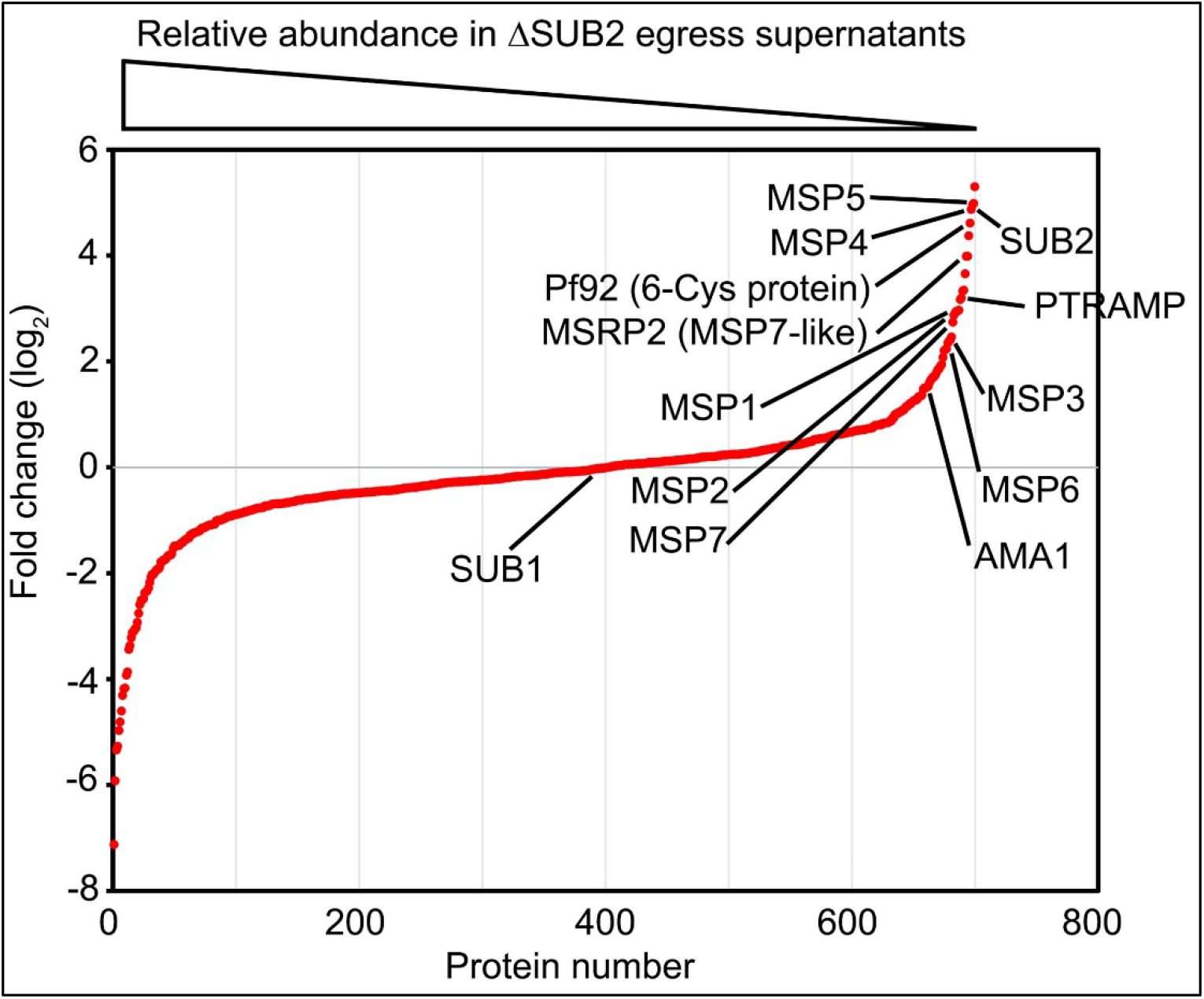
Quantitative proteomics shows that SUB2 sheds multiple merozoite surface proteins. S-curve showing comparative mass spectrometric quantitation of *P. falciparum* proteins in egress supernatants of ΔSUB2 1B4int and control schizonts. A total of 700 parasite proteins were detected with high confidence. Of the top 24 proteins that differed most in abundance between the samples, 12 are known merozoite surface proteins (indicated). SUB1 did not differ in abundance between samples, as expected. See Supplementary Table 1 for mass spectrometric data.

Collectively, these results confirmed SUB2 as the enzyme responsible for shedding of AMA1, PTRAMP and the MSP1 complex as well as several other merozoite surface proteins, and revealed an important role for SUB2 in invasion. The findings also proved that complete shedding of merozoite surface MSP1 and AMA1 is not a prerequisite for invasion as the ΔSUB2 parasites formed rings, albeit with reduced efficiency.

### Confirmation of SUB2 essentiality by genetic complementation; ΔSUB2 parasites that can invade undergo developmental arrest

To explore the long-term consequences of SUB2 loss, ring stage 1B4int parasites were mock-or RAP-treated, dispensed at low density into flat-bottomed microwell plates, and cultured undisturbed. Quantitation of the plaques appearing in the wells (resulting from localised zones of RBC destruction;^36^) revealed that RAP-treated parasites produced significantly fewer plaques than controls (Fig. 4A, B).Genotyping of parasites expanded from 2 of the few plaques in the RAP-treated cultures showed that they were derived from the small fraction of parasites that failed to undergo excision upon RAP treatment (Fig. 4B and Fig. 1E). This suggested that correctly excised parasites lacking an intact *sub2* locus failed to replicate.

**Fig. 4.**
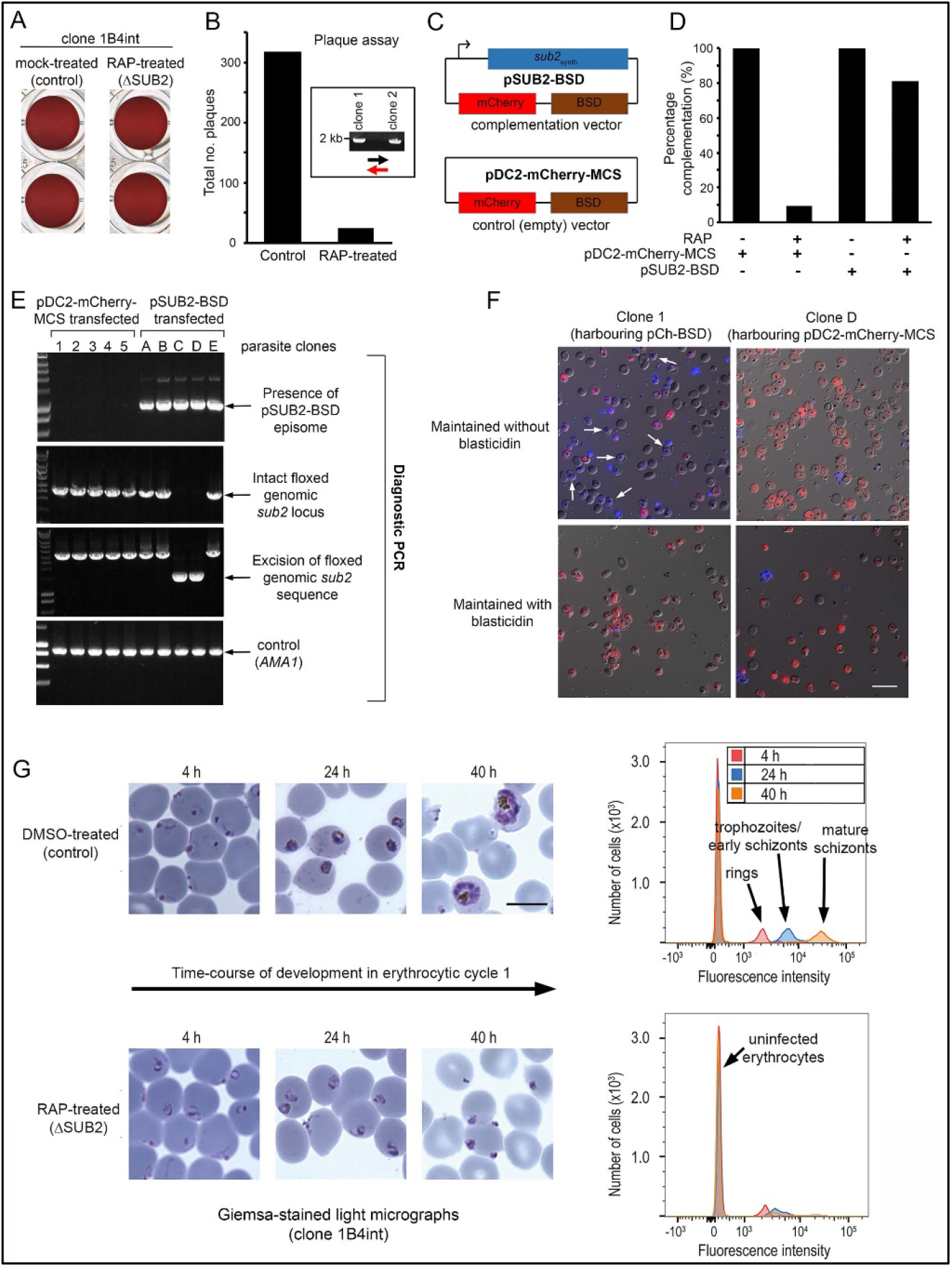
ΔSUB2 parasites display a developmental defect in the erythrocytic cycle following gene disruption that can be rescued by genetic complementation. (A) Plaque assay wells imaged 14 days after plating equal parasite densities (∼10 parasites per well). (B) Plaque numbers in a total of 60 wells per treatment, showing effects of SUB2 disruption. Two plaques from RAP-treated wells were expanded and the parasites confirmed by PCR to be non-excised. See Methods for primer colour-codes. (C) Complementation plasmids. Expression of the *sub2_synth_* gene in pSUB2-BSD was driven by a 5’ flanking region that correctly regulates SUB2 expression ^37^. (D) Genetic complementation of the ΔSUB2 growth defect. Complementation was calculated as the number of plaques obtained in the RAP-treated cultures expressed as a percentage of those in control cultures (100%). (E) Genotyping of parasite clones expanded from the plaque assay in (D). Amplicons diagnostic of indicated genotypes, arrowed. Clones 1-5 (transfected with control plasmid pDC2-mCherry-MCS) and clones A, B and E all possessed an intact genomic *sub2* locus. Clones C and D lacked an intact chromosomal *sub2* gene due to excision, but harboured pSUB2-BSD. (F) Dual DIC/fluorescence images of live Hoechst 33342-stained (blue) parasites of clones 1 and D (from panel (E) following 4 weeks of maintenance. Without BSD, clone 1 began to lose pDC2-mCherry-MCS as indicated by loss of mCherry (red: examples arrowed), whilst pSUB2-BSD was maintained in all clone D parasites. Scale bar, 20 μm. (G) Parasite development in cycle 1 following mock-or RAP-treatment. Arrest of ΔSUB2 parasites was evident within 24 h. Scale bar, 10 μm. Right: flow cytometry confirms a developmental defect in ΔSUB2 parasites. Parasites were stained with SYBR Green I at indicated times post-invasion and 10^5^ cells analysed. See also Supplementary Fig. 2, Supplementary Video 1 and Supplementary Video 2.

To confirm this, and to establish whether the growth defect in the RAP-treated population was solely due to loss of the *sub2* gene, 1B4int parasites were transfected with plasmid pSUB2-BSD designed for ectopic expression of a full-length synthetic SUB2 gene called *sub2*_synth_ ^37^. The plasmid included a blasticidin (BSD) resistance marker and an mCherry expression cassette (Fig. 4C). Parallel 1B4int cultures were transfected instead with a control plasmid (pDC2-mCherry-MCS) lacking the *sub2*_synth_ expression cassette. Following BSD selection, both lines were RAP-treated to disrupt the genomic *sub2* locus, then examined by plaque assay. This showed that carriage of the *sub2*_synth_ transgene, but not the control plasmid, compensated for loss of the genomic *sub2* gene (Fig. 4D). Parasites from single plaques were then expanded in BSD-containing medium and the resulting clones (all mCherry positive) inspected by diagnostic PCR for excision of the floxed chromosomal *sub2* sequence. Whilst none of 25 viable parasite clones harbouring the control plasmid had undergone excision, 2 of 5 selected clones transfected with pSUB2-BSD had undergone excision and so lacked a functional chromosomal *sub2* gene (Fig. 4E). Episomal plasmids segregate inefficiently in *Plasmodium* ^38, 39, 40^ so are usually quickly lost upon removal of drug selection. However, upon further culture of the excised, pSUB2-BSD-harbouring parasites in the absence of BSD, mCherry expression was uniformly maintained (Fig. 4F, top right) indicating that survival was dependent upon maintenance of the pSUB2-BSD expression plasmid. In contrast, the non-excised clones harbouring the control plasmid began to lose mCherry expression in the absence of BSD (Fig. 4F, top left image). These results showed that parasite viability requires a functional *sub2* gene.

To understand the loss of viability in ΔSUB2 parasites, we returned to examine the fate of those ΔSUB2 parasites that successfully invaded at the end of cycle 0. Microscopy and flow cytometry revealed an arrest in intracellular development of the cycle 1 parasites, which - despite appearing initially normal - failed to mature and showed no signs of haemozoin production (a byproduct of haemoglobin digestion) (Fig. 4G and Supplementary Fig. 2). Examination by transmission and serial block-face scanning EM identified no structural defects in the ΔSUB2 cycle 1 rings (Supplementary Fig. 2, Supplementary Video 1 and Supplementary Video 2). We concluded that the lethal phenotype associated with loss of SUB2 arose from a reduced capacity to invade and a developmental arrest in those parasites that did invade.

### ΔSUB2 merozoites lyse target RBCs

During invasion assays involving ΔSUB2 parasites, we noticed that the culture media were unusually red in colour, suggesting a high free haemoglobin (Hb) content. To explore this, mature cycle 0 schizonts of DMSO- or RAP-treated 1B4int parasites were incubated with fresh RBCs for 4 h to allow egress and invasion. Levels of Hb in culture supernatants were then quantified by spectrophotometry and SDS-PAGE, and the cells examined by microscopy and flow cytometry (Fig. 5A). As previously noted, ring production from the ΔSUB2 schizonts was reduced compared to controls. Despite this, higher levels of extracellular Hb appeared in the ΔSUB2 culture supernatants (Fig. 5B). This did not derive from schizont rupture *per se*, as incubation of similar numbers of schizonts without addition of RBCs resulted in low levels of Hb release that did not differ between ΔSUB2 and control schizonts. Moreover, Hb release required extensive interaction between released merozoites and the host cells, since it did not occur in the presence of cytochalasin D (cytD), an actin-binding drug that blocks invasion downstream of TJ formation by disrupting the parasite actinomyosin motor that drives invasion ^41^ (Fig. 5C). We conjectured that those ΔSUB2 merozoites unable to invade instead interacted with target RBCs in an abortive manner that led to lysis. To investigate this, following co-incubation of schizonts with RBCs, cultures were supplemented with fluorescent phalloidin, a membrane-impermeable peptide that binds the F-actin of the RBC cytoskeleton ^42, 43^, then immediately examined by live fluorescence microscopy. This revealed rounded, phalloidin-labelled cells that were more abundant in the ΔSUB2 cultures (Fig. 5D). The dimensions of these cells, their low buoyant density (shown by a tendency to ‘float’ above intact RBCs) and their accessibility to phalloidin, suggested that they were ‘ghosts’ derived from lysis of RBCs. We concluded that these derived from abortive invasion attempts in which interaction with ΔSUB2 merozoites had induced RBC lysis.

**Fig. 5.**
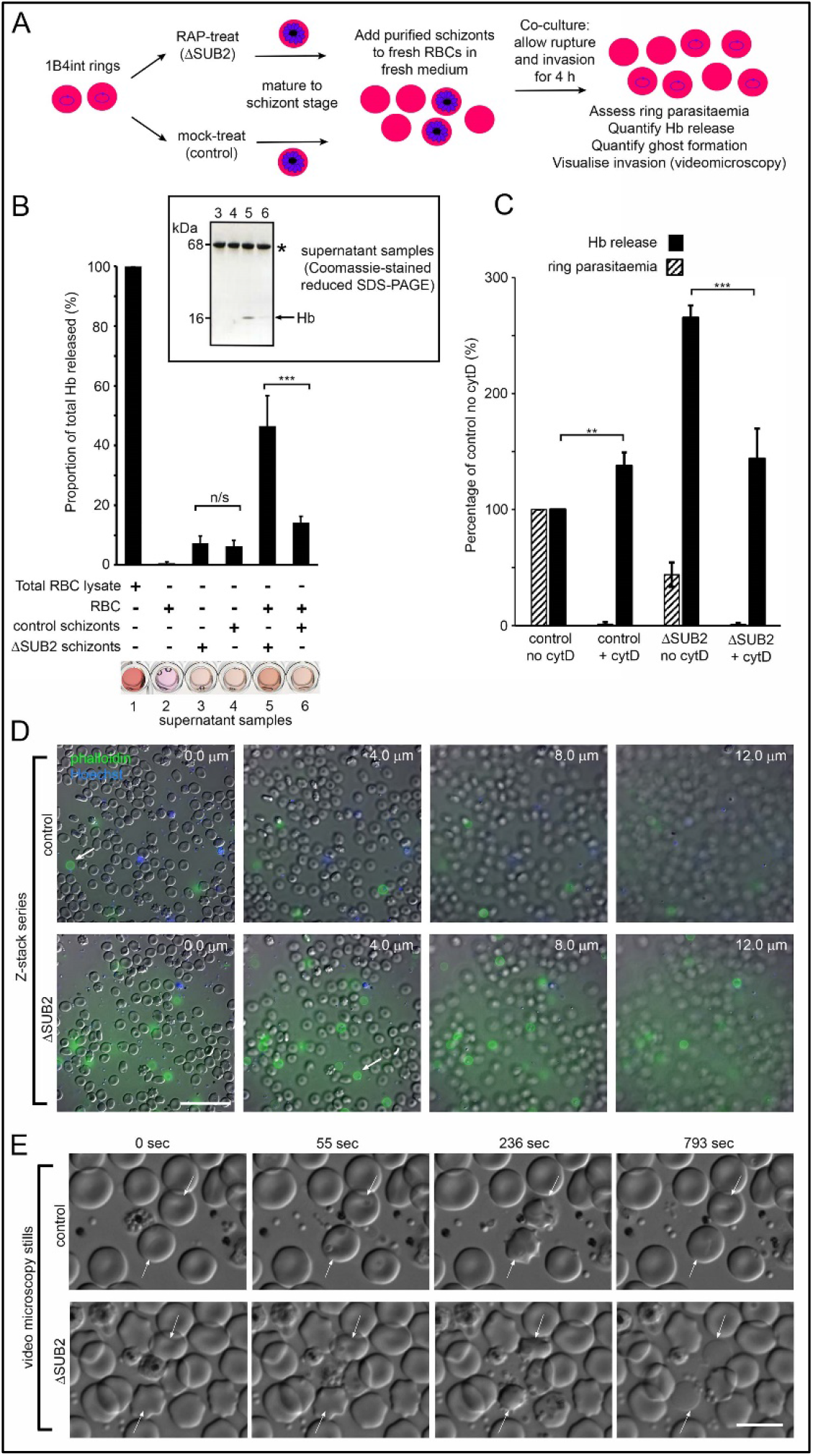
Interaction of ΔSUB2 merozoites with target RBCs induces rapid RBC lysis. (A) Experimental strategy to quantify the relationship between invasion and Hb release. (B) Quantitation of Hb in culture supernatants following egress and invasion. Values shown (averages from 3 independent experiments) were determined by absorbance at 405 nm, normalised to hypotonic lysates of an equivalent number of uninfected RBCs (100%; see Methods). These experiments used high starting schizont parasitaemia (10-15%) to enhance detection of released Hb. Mean ring parasitaemia values following 4 h of ΔSUB2 schizont/RBC co-culture were only 65.0 ±4.8% of those in the control schizont/RBC co-cultures, whereas released Hb levels in the ΔSUB2/RBC co-cultures were 3.3 ±0.7-fold higher than in the control schizont/RBC co-cultures (***). Error bars, ±SD. Significance determined by two-tailed unpaired t-test: p = 0.006, ***; n/s, p = 0.560. Below, microplate wells containing culture supernatants (200μl) compared to the RBC lysate, showing the high free Hb levels in the ΔSUB2 schizont/RBC co-cultures. Inset: SDS-PAGE analysis of supernatants 3-6. (C) Hb release is prevented by cytD. Ring parasitaemia and released Hb levels relative to those obtained in co-cultures of control schizonts in the absence of cytD (defined as 100%). While Hb release is enhanced throughout by cytD, the drug blocks the high levels of Hb release observed in the ΔSUB2 co-cultures. Shown are averages from 2 independent experiments. Error bars, ±SD. Significance levels determined by two-tailed unpaired t-test: p = 0.03, ***; p = 0.04, **. (D) Interaction of ΔSUB2 merozoites with RBCs leads to abundant ghost formation. Cultures as in (A) were supplemented with Hoechst 33342 (blue) and Alexa Fluor 488 phalloidin (green) then imaged by dual DIC/fluorescence microscopy. Images are selected from a Z-stack series of single fields, taken at 2.0 μm intervals to detect phalloidin-labelled RBC ghosts, most of which ‘float above’ the focal plane of the majority of intact RBCs. Ghosts were ∼6-fold more numerous in the ΔSUB2 cultures (15.7% of the total RBC count in ΔSUB2 cultures; 2.6% in control cultures). Scale bar, 50 μm. (E) Abortive invasion by ΔSUB2 merozoites leads to RBC lysis. Images from live time-lapse DIC microscopy of egress and subsequent events. Time following start of imaging is indicated. Interaction with target RBCs (arrowed) results in transient echinocytosis, followed by lysis in the case of the ΔSUB2 parasites. Scale bar, 10 μm. See also Supplementary Video 3.

To directly visualise the fate of target RBCs, we examined the process by live microscopy (Fig. 5E and Supplementary Video 3). Initial interactions with ΔSUB2 merozoites appeared normal, with RBC deformation often followed by parasite entry and echinocytosis. Subsequently however, and in contrast to the behaviour typical of RBCs invaded by control merozoites where echinocytosis quickly resolved, RBCs invaded by ΔSUB2 merozoites often remained rounded and then lysed, as indicated by loss of differential interference contrast (DIC). Lysis often occurred within ∼14 minutes of the initial interaction and was sometimes accompanied by ejection of the merozoite. In other cases lysis took longer and was visualised by images captured over 30-60 minutes. Attempts to fix ΔSUB2 merozoites in the act of invasion for analysis by EM were unsuccessful, possibly due to its transient nature. Nonetheless we concluded that loss of SUB2 led in ∼50% of invasion attempts to abortive invasion that culminated in RBC lysis, presumably due to a defect in RBC sealing.

### Inhibition of MSP1 shedding by cleavage site mutagenesis inhibits intracellular parasite development

To dissect the ΔSUB2 phenotype, we investigated the effects of mutations that directly prevent SUB2-mediated shedding of the most abundant SUB2 substrate, the MSP1 complex. We previously mapped the SUB2 cleavage site in MSP1 to the Leu1606-Asn1607 bond just upstream of the C-terminal MSP1_19_ domain ^12^ and showed that proline substitutions of the residues flanking the AMA1 SUB2 cleavage site blocks cleavage ^25^. Based on this, transgenic parasite line iMSP1_PP_ was generated in which DiCre-mediated excision of a floxed segment of the *msp1* gene produced a partial allelic replacement, substituting the Leu-Asn cleavage motif with a Pro-Pro motif (Supplementary Fig. 3). In a control parasite line, iMSP1_LN_, excision reconstituted the wild type motif.

RAP-treatment of iMSP1_PP_ and iMSP1_LN_ parasite clones produced the expected genomic excision events (Supplementary Fig. 3). To examine the effects of the Leu-Asn to Pro-Pro substitution, synchronous rings were RAP- or mock-treated, matured to schizont stage, then allowed to undergo egress in the presence of RBCs. This showed no significant effects on efficiency of ring formation (Fig. 6A), although examination of culture supernatants following invasion showed a selective reduction in MSP1 shedding in RAP-treated iMSP1_PP_ cultures, consistent with the predicted effects of the mutations on SUB2-mediated cleavage (Fig. 6B). Cycle 1 rings from RAP-treated iMSP1_PP_ schizonts were strongly recognised by mAb X509, similar to ΔSUB2 cycle 1 rings (Fig. 6C), indicating successful invasion despite reduced MSP1 shedding. The newly-invaded cycle 1 mutant iMSP1_PP_ rings appeared morphologically normal, but further monitoring revealed extensive retarded intracellular development (Fig. 6D), and longer-term experiments confirmed a replication defect in the RAP-treated iMSP1_PP_ parasites (Fig. 6E). We concluded that selective inhibition of MSP1 shedding at invasion did not affect RBC entry but led to defective intracellular parasite development resembling that in the ΔSUB2 mutants.

**Fig. 6.**
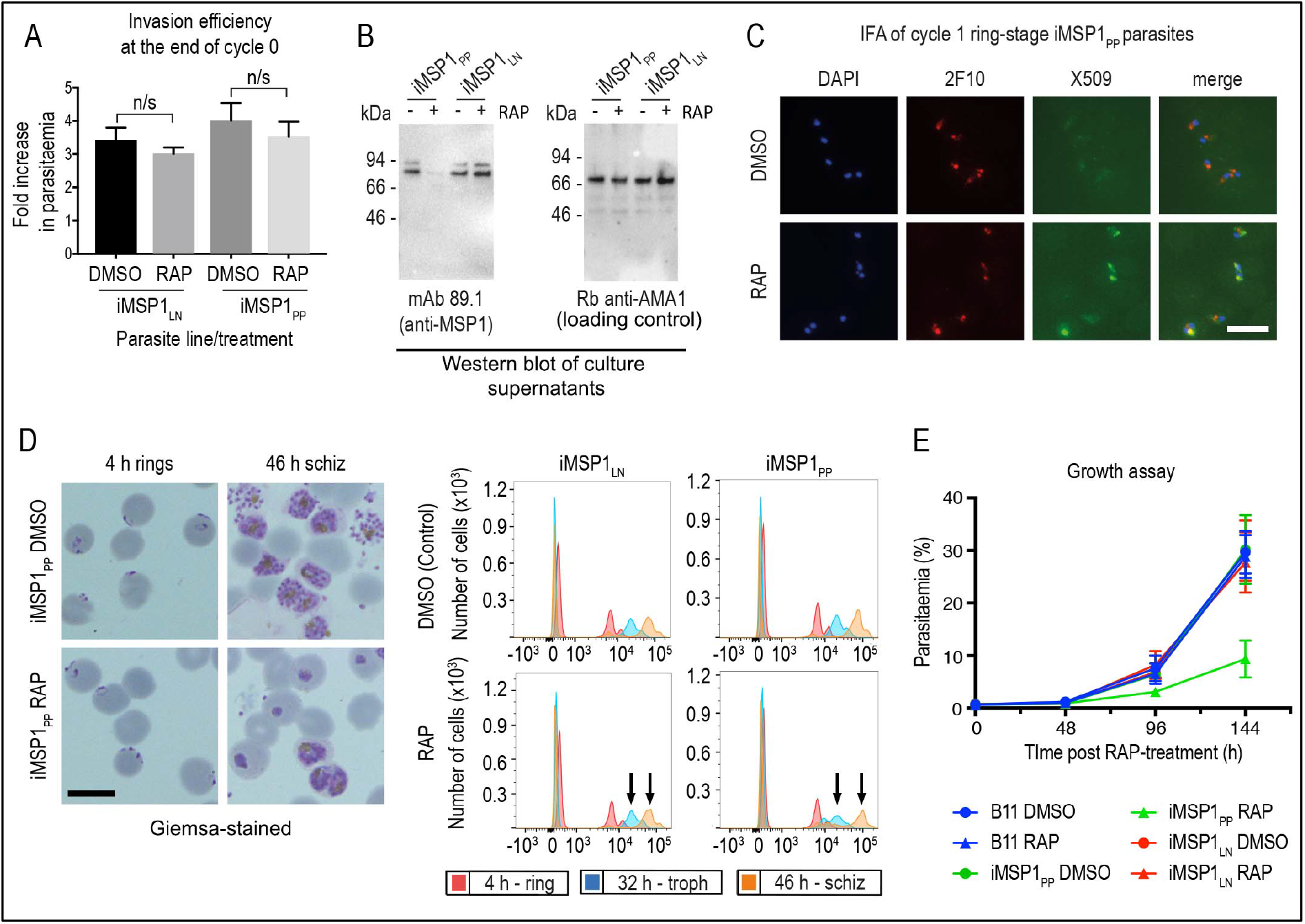
Inhibition of MSP1 shedding by conditional mutagenesis of the SUB2 cleavage site inhibits cycle 1 intracellular parasite development. (A) Ring formation is unaffected by conditional mutagenesis of the MSP1 SUB2 processing site. Values averaged from 3 independent experiments. Error bars, ±SD. Significance determined by two-tailed t-test: p values for the comparison of DMSO and RAP-treated MSP1_LN_ and MSP1_PP_ lines were 0.1662 and 0.2988 respectively (n/s). (B) Reduced MSP1 shedding during invasion by RAP-treated iMSP1_PP_ parasites. (C) IFA showing that cycle 1 rings formed by RAP-treated iMSP1_PP_ parasites were recognized by mAb X509, consistent with reduced MSP1 shedding. All rings are detected by the MSP1_19_-specific mAb 2F10. (D) Reduced MSP1 shedding affects intracellular development. Control or RAP-treated iMSP1_LN_ and iMSP1_PP_ schizonts were incubated with RBCs and the cycle 1 rings monitored. Microscopy shows retarded ring development in RAP-treated iMSP1_PP_ parasites, corroborated by flow cytometry (measuring parasite DNA content) showing 23.6 ± 5.0% reduced replication by 46 h compared to similarly treated iMSP1_LN_ controls (compare arrowed peaks). Scale bars, 10 μm. E) Reduced replication of RAP-treated iMSP1_PP_ parasites. See also Supplementary Fig. 3.

### Inhibition of AMA1 shedding by cleavage site mutagenesis has no impact on invasion but loss of AMA1 expression prevents invasion and results in host RBC lysis

The MSP1 mutagenesis data provided a plausible explanation for the impact of SUB2 loss on intracellular parasite growth, but did not explain the invasion phenotype of ΔSUB2 merozoites nor their capacity to lyse RBCs. Prior to this work, the only other experimentally-demonstrated essential SUB2 substrate was the microneme protein AMA1. The AMA1 ectodomain comprises three globular domains linked to the TMD via a short juxtamembrane segment ^44^. Cleavage of *P. falciparum* AMA1 by SUB2 occurs at the Thr517-Ser518 bond 29 residues upstream of the TMD, releasing domains I-III, leaving a juxtamembrane ‘stub’ bound to the merozoite surface ^17^. AMA1 can additionally be shed by the parasite rhomboid protease ROM4 via cleavage within the TMD at the Ala550-Ser551 bond ^17, 29^; this can be blocked by a Tyr substitution of Ala550, a mutation tolerated by the parasite ^25^. To probe the biological significance of SUB2-mediated shedding of AMA1, we conditionally modified both the SUB2 and ROM4 cleavage sites to simultaneously render them refractory to cleavage. For this, we generated transgenic parasite line iΔRΔS in which excision of a floxed *ama1* gene simultaneously substitutes the Thr517-Ser518 site with a Pro-Pro motif and Ala550 with a Tyr residue (Supplementary Fig. 4).

Replication of untreated iΔRΔS parasites was indistinguishable from parental parasites, and RAP-treatment produced the expected genomic changes (Supplementary Fig. 4) as well as the anticipated decrease in AMA1 shedding (Fig. 7A). However, we detected no change in the invasive capacity of the RAP-treated parasites and replication was unaffected (Fig. 7B). Given the absence of deleterious effects, we reasoned that direct mutagenesis of the AMA1 cleavage sites should generate viable parasites, so we used targeted homologous recombination to directly introduce the same cleavage site mutations into the *ama1* gene (Supplementary Fig. 4). The resulting parasite line, dΔRΔS, displayed a phenotype similar to that of the RAP-treated iΔRΔS parasites, with reduced AMA1 shedding (Fig. 7A) but no effects on invasion or replication (Fig. 7B). These results suggested that inhibition of AMA1 shedding *per se* was not responsible for the invasion and lysis phenotype observed in the ΔSUB2 mutants.

**Fig. 7.**
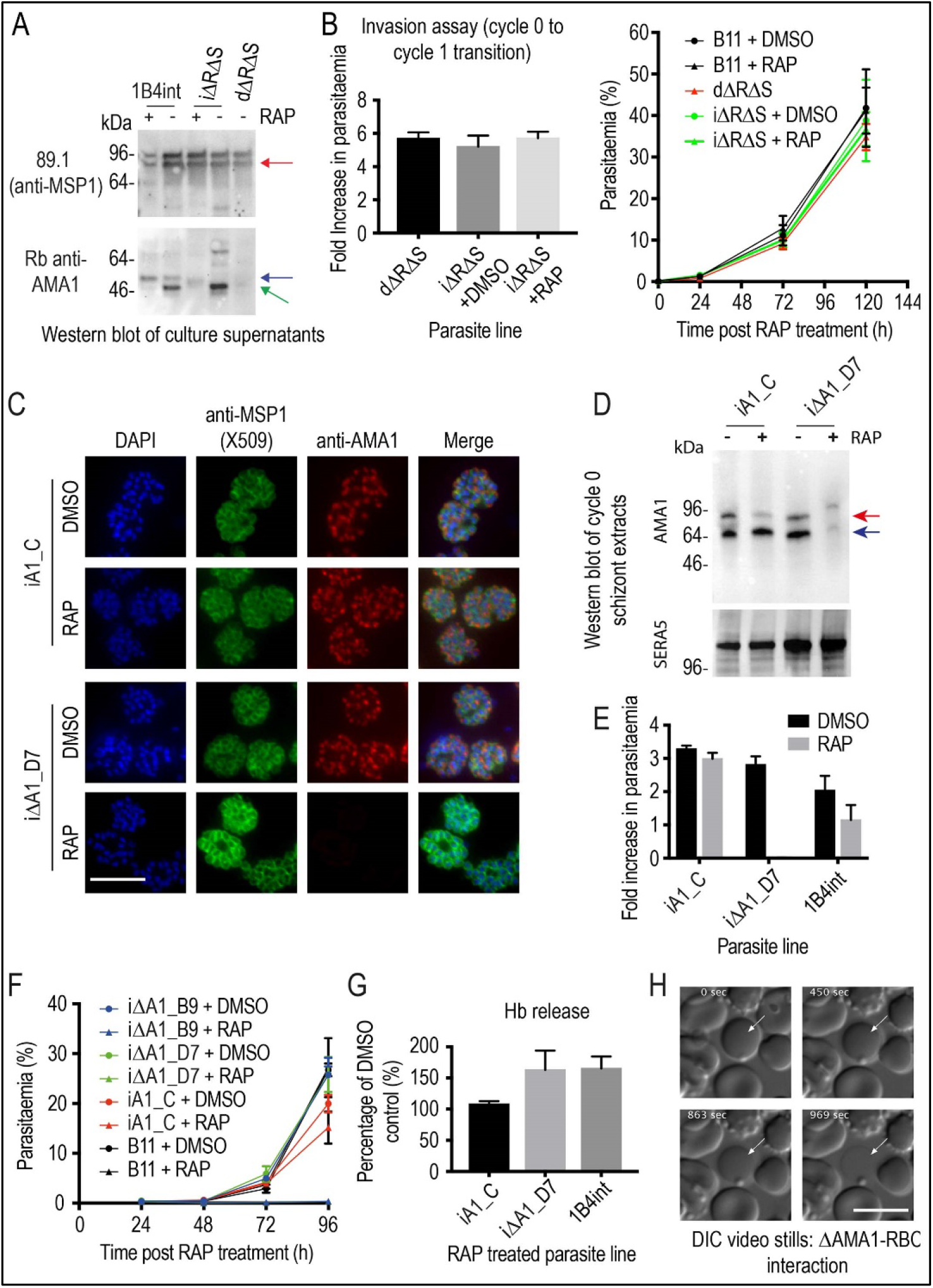
Inhibition of AMA1 shedding by cleavage site mutagenesis has no effect on invasion, but AMA1 disruption prevents invasion and leads to RBC lysis. (A) Selective reduction in AMA1 shedding in dΔRΔS and RAP-treated iΔRΔS parasites. Shed fragments are arrowed (MSP1_83_, red; AMA1_52_, blue; AMA1_48_, green). (B) Cleavage site mutagenesis has no impact on invasion or parasite replication. Data in both cases are averages of 3 independent experiments. Error bars, ±SD. (C) IFA showing loss of AMA1 expression in RAP-treated iΔAMA1 cycle 0 schizonts. (D) Disruption of AMA1 expression (arrowed) in RAP-treated iΔAMA1 parasites. Antibodies to SERA5 were used as a loading control. (E) Invasion assay showing complete loss of invasion following rupture of RAP-treated cycle 0 iΔAMA1 schizonts in the presence of RBCs. (F) Loss of replication of RAP-treated iΔAMA1 parasite clones. (G) Increased Hb release upon interaction of ΔAMA1 parasites with RBCs is similar to that in RAP-treated 1B4int (ΔSUB2) parasites. (H) Time-lapse DIC microscopy images of interactions between a ΔAMA1 merozoite and an RBC (arrowed), showing eventual lysis of the RBC. Time following start of imaging is indicated. Scale bars, 10 μm. See also Supplementary Fig. 4, Supplementary Fig. 5 and Supplementary Video 4.

Given the role of AMA1 in TJ formation, we examined whether the ΔSUB2 phenotype might in part reflect loss of AMA1 function. We generated a parasite line (iΔAMA1) in which DiCre-mediated excision ablated AMA1 expression by severely truncating the gene (Supplementary Fig. 5). In a control line (iAMA1_C), excision reconstituted a functional full-length gene (Supplementary Fig. 5). RAP-treatment of the parasites produced the expected genomic modifications (Supplementary Fig. 5), with loss of AMA1 expression in mature cycle 0 schizonts of the iΔAMA1 clones (Fig. 7C, D). This led to complete loss of ring formation and proliferation following rupture of RAP-treated iΔAMA1 schizonts at the end of cycle 0 (Fig. 7E, F), confirming AMA1 essentiality. Despite the lack of invasion, levels of Hb released into the culture media were much higher in RAP-treated iΔAMA1 schizonts incubated with RBCs than in controls (Fig. 7G). To determine the source of the Hb, we used microscopy to visualise interactions between iΔAMA1 merozoites and RBCs. This confirmed the loss of invasion, whilst showing that interaction of ΔAMA1 merozoites with target RBCs produced extensive RBC echinocytosis (Supplementary Video 4). We did not observe rapid RBC lysis; however, target RBCs often transformed to a rounded form, remaining in this state for some time before losing Hb content (Fig. 7H). We concluded that loss of AMA1 not only prevented invasion but also produced host RBC lysis upon merozoite interaction, presumably due to an RBC sealing defect similar to that associated with loss of SUB2.

## Discussion

We have provided the first genetic proof of SUB2 as the merozoite sheddase, and have shown that abolishing SUB2-dependent cleavage produces a complex phenotype that is lethal in at least two superficially distinct manners; merozoites that successfully complete invasion fail to develop, whilst other merozoites induce RBC lysis at or shortly following entry. Our most important conclusion is that SUB2-mediated protein shedding is essential for parasite survival. Our primary mechanistic explanation for this is that shedding is required for resealing of the RBC membrane at invasion, but credible additional factors are worth considering.

Using IFA, Western blot and mass spectrometry to compare SUB2-expressing and ΔSUB2 parasite cultures, we confirmed that SUB2 is required for shedding of MSP1, AMA1 and PTRAMP. We also identified several putative new SUB2 substrates not known to associate with any of the three previously known substrates. Some of the most prominent of these, including MSP2, MSP4, MSP5 and Pf92, belong to a group of merozoite surface proteins known or predicted to possess GPI anchors ^30, 45, 46, 47^. Since GPI-anchored proteins cannot be shed by rhomboid cleavage (which occurs within TMDs), it is likely that shedding of each is independently catalysed by SUB2. The other newly identified SUB2 substrate, MSRP2, likely has no GPI anchor (so could be peripherally associated with one of the GPI-anchored substrates) but does undergo proteolytic cleavage around egress ^48^. This wide range of SUB2 substrates was unexpected; indeed earlier work that did not benefit from access to mutants concluded that MSP2 and MSP4 are not shed at invasion ^49^. Our new findings underline the potential for multiple consequences of SUB2 inhibition.

Despite the reduction in merozoite surface protein shedding, ΔSUB2 parasites formed new rings at the end of cycle 0, albeit at ∼50% reduced efficiency. These uniformly arrested, and growth assays supported by genetic complementation proved that SUB2 is indispensable for parasite survival. To dissect the phenotype, we generated lines in which shedding of the most abundant SUB2 substrate, the MSP1 complex, was selectively inhibited. These mutants invaded with wild type efficiency but displayed a developmental defect similar to that of the ΔSUB2 mutants. Given the technical difficulties in determining causality upon intracellular arrest in *Plasmodium*, we can only speculate on the underlying defect(s). The MSP1_19_ species remaining on the merozoite surface following SUB2-mediated shedding has been implicated in formation of the acidic digestive vacuole (DV) that is the site of haemoglobin catabolism during intraerythrocytic development ^50^. We speculate that in the absence of shedding, the bulky MSP1 complex remaining on the plasma membrane of the internalised parasite, likely together with other unshed SUB2 substrates, interferes with DV biogenesis, inhibiting haemoglobin digestion and stalling growth. The lack of haemozoin in both the cycle 1 ΔSUB2 parasites and many arrested iMSP1_PP_ mutants is consistent with this model.

The most dramatic phenotype associated with loss of SUB2 was extensive lysis of targeted RBCs. What could underlie this? The mechanisms regulating PVM formation, resealing and scission during invasion by apicomplexan parasites are not understood. Numerous EM studies (e.g. ^1^ indicate that, topologically, invasion resembles induced invagination of the RBC membrane, so that the host RBC cytosol is never exposed to the extracellular milieu. Conflicting with this is recent evidence that successful RBC invasion by *P. falciparum* merozoites involves the formation of a transient discontinuity in the host cell membrane, allowing Ca^2+^ flux into the RBC ^3, 51^. Similarly, an elegant time-resolved patch-clamp study in the related apicomplexan *Toxoplasma gondii* detected a spike in host cell membrane conductance upon apical attachment of the parasite, again consistent with pore formation^52^. Near the end of invasion, as the parasite posterior enters the nascent PVM, it seems likely that a membrane scission event seals the shrinking TJ aperture, and evidence for this too was found in the *Toxoplasma* study ^52^. Recent work in *Toxoplasma* suggests that PVM scission is aided by rotation of the parasite along its apical-posterior axis immediately following entry ^53^, and malaria merozoites have also been observed to spin post invasion ^2, 54^.

Given these models of pore formation early in invasion and a membrane scission event at the end of the process, it seems plausible that a defect in either could cause a localised loss of host cell membrane integrity sufficiently catastrophic to result in lysis. RBC lysis associated with defective or delayed entry has been previously documented even in wild type *Plasmodium* ^2, 54^, and indeed we noticed phalloidin-labelled RBC ghosts at low frequency in our control cultures, so the process is clearly sensitive to perturbation. We suggest that RBC lysis associated with loss of SUB2 is at least in part due to an accumulation of unshed merozoite surface proteins at the narrowing TJ aperture, preventing sealing. That lysis does not occur in the presence of cytD, which blocks invasion at the point of TJ assembly ^41^, is consistent with this. Our MSP1 mutagenesis results show that inhibition of MSP1 shedding is not alone sufficient to cause lysis, but since MSP1 is just one of many SUB2 substrates this mutant does not accurately reflect the ΔSUB2 phenotype.

The ΔSUB2 sealing defect may be magnified by the fact that the SUB2 substrate AMA1 is a core component of the TJ, bridging the parasite to the RBC through interactions with RON2, a member of a complex of rhoptry neck proteins inserted into the host cell membrane early in the invasion pathway, perhaps at pore formation ^55, 56, 57, 58, 59, 60^. The AMA1-RON2 interaction may play a role in drawing the TJ orifice closed as the parasite posterior enters the PV. If so, an attractive concept is that SUB2-mediated release of the AMA1 ectodomain may facilitate pinching off of the PVM, and so we initially considered that loss of AMA1 shedding might be primarily responsible for the ΔSUB2 lysis phenotype. We were therefore surprised to find that selective inhibition of AMA1 shedding had no impact on invasion or viability. In contrast, disruption of AMA1 expression *did* produce a lysis phenotype, as well as the expected loss of invasion. We reconcile these disparate observations by speculating that lysis occurs upon loss of AMA1 because the RBC membrane is compromised early in invasion by insertion of the RON complex (which occurs independently of the presence of AMA1; ^59, 61^) but subsequent stabilisation of the TJ cannot occur. While reducing AMA1 shedding by cleavage site mutagenesis does not in isolation prevent sealing, failure to shed AMA1 in combination with failure to shed multiple other proteins – as in the case of the ΔSUB2 mutant – leads to a profound sealing defect and rapid RBC lysis.

A role for *P. falciparum* AMA1 in RBC sealing has been postulated previously, based on work using a less efficient conditional mutagenesis strategy which led to variable levels of AMA1 depletion^62^. In that study many parasites that invaded failed to develop, leading the authors to suggest that even a partial sealing defect could stall development. We concur, and although we detected no membrane defects in the ΔSUB2 cycle 1 rings, the developmental arrest we observed in the ΔSUB2 and MSP1 cleavage site mutants might also stem from a sealing defect insufficiently severe to cause lysis. Interestingly, neither the Yap et al. study nor other studies of *Plasmodium* or *Toxoplasma* AMA1-null mutants noted widespread host cell lysis ^61, 63, 64^. We suspect three likely reasons for this. First, our conditional gene disruption system is highly efficient, with close to 100% conversion in a single erythrocytic cycle. Second, in *Toxoplasma* disruption of AMA1 can lead to upregulation of AMA1 paralogues absent in *Plasmodium* ^59, 65^. Third, RBCs are more sensitive to lysis resulting from membrane wounding than the nucleated cells invaded by *Toxoplasma*, perhaps due to the absence in RBCs of endomembrane-dependent repair mechanisms ^66^.

In summary, the lethal consequences of SUB2 disruption likely result from: (1) a combination of loss of shedding of multiple surface proteins plus interference with AMA1 function, leading to defective RBC sealing; and (2) arrested post-invasion development due to inhibition of DV biogenesis and/or incorrect sealing. Our work identifies SUB2 as a critical mediator of host RBC integrity and parasite viability. Drug-like inhibitors of SUB2 protease activity have potential as a new class of antimalarial drug that would inhibit invasion and parasite replication.

## Supporting information

Vidoe S3

Vidoe S4

## Acknowledgements

This work was supported by funding to MJB from the Francis Crick Institute (https://www.crick.ac.uk/) which receives its core funding from Cancer Research UK (FC001043; https://www.cancerresearchuk.org), the UK Medical Research Council (FC001043; https://www.mrc.ac.uk/), and the Wellcome Trust (FC001043; https://wellcome.ac.uk/). The work was also supported by Wellcome ISSF2 funding to the London School of Hygiene & Tropical Medicine.

## Author Contributions

CRC, FH, MJB; Conceptualization. CRC, FH, MJB, SAH, APS, MRGR; Formal analysis. APS, LMC, MJB; Funding acquisition. CRC, FH, SAH, MRGR, MJB; Investigation. APS, LMC, MJB; Supervision. CRC, FH, MJB; Writing – original draft. CRC, FH, APS, MJB; Writing – review and editing.

## Methods

### Materials and *P. falciparum* culture

See Supplementary Table 2 for a list of all reagents, cell lines and software used in this study. Asexual blood-stages of *P. falciparum* were maintained at 5-10% parasitaemia in RPMI1640 supplemented with 0.5% Albumax II (RPMI 1640-Albumax; ThermoFisher) (Trager and Jensen,1976). Parasites were synchronized at 48-96 h intervals using standard methodology^74^. Briefly, mature schizonts were enriched by centrifugation over a 70% isotonic Percoll cushion (GE Healthcare Life Sciences) and then allowed to invade fresh AB+ RBCs (NHSBT) for 1-2 h. Following invasion, remaining intact schizonts and schizont debris were removed by centrifugation over a 70% isotonic Percoll cushion and the newly-invaded ring stages further treated with 5% (w/v) sorbitol for 7 min at 37°C to lyse any residual schizonts. The final ring cultures were washed and returned to culture or used as required.

### Construction of plasmid constructs to genetically modify *P. falciparum*

PCR amplicons used in plasmid cloning were generated using Fusion high fidelity DNA polymerase (ThermoFIsher) or Platinum Taq DNA polymerase, High Fidelity (ThermoFisher) and purified using Qiagen PCR purification or Qiagen Gel extraction kits.

#### Construct for generation of conditional SUB2 disruption parasite clones 1B4int and FE5int

A fragment of *sub2* sequence (S2syn) was commercially synthesised (GeneArt), comprising a stretch of native *P. falciparum* 3D7 *sub2* sequence followed by the 3D7 *sub2* intron containing an internal *loxP* site (*loxPS2Int*) and finally a stretch of recodonised *sub2* gene sequence encoding the 3D7 amino acid sequence but using a different codon usage ^14^. The 3’ end of the recodonised *sub2* gene was amplified from plasmid psub2-sub2wHA3_F1 ^37^ using primers sgS2_HBA_F and sgS2_X1_R and cloned into plasmid pHH1_sera5_LoxP1 ^28^ which has a *loxP* site downstream of the Xho I site at the 3’ end of the *sera5* gene sequence, using Hpa I and Xho I, generating plasmid pHH1_sgS2_3HA. A *sub2* target region was amplified from 3D7 genomic DNA using primers S2endo_HpaI_F and S2endo_BspEI_R and cloned into pHH1_sgS2_3HA using Hpa I and Bsp EI resulting in plasmid pHH1_endo/sgS2_3HA. Plasmid pHH1_sub2_KO was obtained by cloning S2syn into pHH1_endo/sgS2_3HA using Bsp EI and Age I. The unmodified endogenous and modified (integrated) *sub2* loci were detected by PCR using forward primer SUB2for_Int and reverse primers SUB2rev_wt and SUB2rev_Int (1.95 kb) respectively. The intact and excised *sub2* loci following RAP-induced recombination were detected with forward primers S2assplasfor2 and nonexSUB2_F respectively and reverse primer pHH1R1 (giving rise to 1.7 and 2.6 kb fragments respectively).

#### SUB2 complementation plasmid

The *sub2* expression cassette was excised from plasmid pSUB2_sub2wHA_PEX ^37^ using Not I and Spa I. This sequence was cloned into pDC2-mCherry-MCS ^75^ pre-digested with Not I and Hpa I giving rise to plasmid pSUB2-BSD. Plasmid pDC2-mCherry-MCS was used as the control plasmid for complementation. The presence of episome was detected by PCR using primers sgSUB2_5 and sgSUB2_31 (amplicon size; 1.0 kb). PCR amplification of the *ama1* gene locus with primers A1seq1for and A1seq6rev was used as a loading control (0.9 kb).

#### Construct design to generate transgenic parasite lines iMSP1_PP_ and iMSP1_LN_ for conditional expression of mutant MSP1

Three synthetic gene fragments were generated (GeneArt). These were: (1) Fragment A: chimera of the 3D7 *msp1* sequence and Wellcome *msp1_19_* with an internal *loxPint* sequence. This sequence was flanked by Bst BI and Xho I restriction sites for cloning purposes; (2) Fragment B: *loxPint* followed by the 3D7 *msp1_19_* sequence and endogenous 3D7 *msp1* 3’ UTR. The sequence encoding the SUB2 cleavage site (Leu-Asn) in the 3D7 *msp1* sequence was mutated to encode Pro-Pro. The synthetic fragment was flanked by Not I and Nde I restriction sites; (3) Fragment C: *loxPint* followed by 3D7 *msp1_19_* sequence. A chimeric sequence comprising 3D7 target, Wellcome *msp1_19_* and PbDT3’ UTR was excised from pHH1-wMSP1-wt ^76^ using Hpa I and Not I and cloned into pGEM®-T Easy pre-digested with Sph I (T4 polymerase blunt ended) and Not I, generating plasmid pGEM-MSP1. Fragment A was cloned into pGEM-MSP1 using Xho I and Bst BI giving rise to plasmid pGEM-MSP1(A). Plasmid pGEM-MSP1_PP_ was generated by cloning B into pGEM-MSP1(A) using Not I and Nde I restriction digestion. Control plasmid pGEM-T MSP1_LN_ was obtained by cloning fragment C into plasmid pGEM-MSP1_PP_ using Not I and Pst I. All PCR screening relied on forward primer MSP1_Int_F in combination with MSP1_EndCont_R for the endogenous locus (amplicon: 1.4 kb) and wMSP119_Int_R for integrated (amplicon: 1.4 kb) and excised (amplicon: 1.4 kb) loci.

#### Construct design to generate parasites conditionally expressing AMA1 refractory to cleavage by SUB2 and ROM4 (iΔRΔS)

A synthetic sequence was ordered comprising a small section of the final 3D7 *ama1* target region, followed by recodonised segment of the *ama1* 3’ from *P. falciparum* strain FVO^77^ flanked by *loxPS2int* sequence, 3D7 *ama1* 3’ and finally native 3D7 *ama1* 3’ UTR (GeneArt). The recodonised FVO sequence was modified by amino acid replacement to include a haemagglutinin (HA) epitope tag within the stub region between domain III and the transmembrane region of AMA1 (IPEHKPTYD/ YPYDVPDYA) as previously described ^78^. The native *ama1* 3’ sequence was altered to introduce the TS/PP mutations at the P1-P1’ positions of the SUB2 cleavage site, as well as an HA epitope tag. This HA tag was introduced by amino acid replacement into an unstructured loop in domain III (KRIKLNDND/YPYDVPDYA) as previously described ^60^. Native 3D7 *ama1* target sequence was amplified from 3D7 genomic DNA using primers endo_SacII_F and endo_dsBclI_R and cloned into pGEM-T using SacII and NdeI, giving rise to plasmid pGEM-pfama1. The GeneArt generated synthetic sequence was cloned into pGEM-pfama1 using Sac II and Nde I generating plasmid pGEM-A1-3’UTR. The PbDT3’ sequence was excised from pHH1-5’sgPfa1HA ^78^ and inserted between the FVO *ama1* sequence and the second *loxPS2int* using Avr II and Not I giving rise to pGEM-T-iΔS_A1.

The region encoding the SUB2 cleavage site was amplified from plasmid pGEM-T-iΔS_A1 using primer pairs endo_usPacI_F/A1_S2_C_R and A1_S2_C_F/A1_dsBsu361_R. Overlapping PCR was carried out using the resulting fragments and primers endo_usPacI_F and A1_dsBsu361_R. The resulting PCR product was cloned into plasmid pGEM-T-iΔS_A1 using restriction enzymes PacI and Bsu361 giving rise to the control plasmid pGEM-T-iA1_C.

Mutations at the P1 position of the rhomboid cleavage site (Y550A) in the region encoding the AMA1 TMD were introduced by overlapping PCR. Briefly, PCR fragments were obtained from pGEM-T-iΔS_A1 using primer pairs endo_usPac1/A1_A550_Y_R and A1_A550_Y_F/ama1 3’_dsNde1. Overlapping PCR was carried out using primers endo_usPac1 and ama1 3’_dsNde1. This product was cloned into pGEM-T-iA1_C and pGEM-T-iΔS_A1 giving rise to plasmids pGEM-T-iΔR_A1 and pGEM-T-iΔRΔS_A1. PCR screening relied on use of the forward primer A1_Int_F1 in combination with reverse primers A1_EndCont_R1 for the endogenous locus (amplicon; 1.3 kb), A1_Int_R for the integrated locus (1.4 kb) and A1_dsBsu361_R for both non-excised (3.9 kb) and excised (1.9 kb) loci.

#### Construct design for direct insertion of SUB2/ROM4 cleavage site mutations in the ama1 gene (line dΔRΔS)

Two gBlocks were ordered (IDT) comprising recodonised *ama1* sequence from upstream of the Avr II site and incorporating a 3’ Bsu361 site at the position equivalent to that in the endogenous 3D7 sequence. The control gBlock (C) had no additional modifications while the other had mutations at the SUB2 and rhomboid cleavage sites described above (ΔRΔS). The gBlocks were cloned into pCR™-Blunt (Thermo Fisher Scientific) and then excised using Avr II and Bsu361 and cloned into pGEM-T-iA1-C pre-digested with the same restriction enzymes giving rise to plasmids pGEM-T_dA1_C and pGEM-T_dA1_ΔRΔS. PCR screening was the same as for the conditionally uncleavable AMA1 lines (iΔRΔS_A1).

#### Construct for generation of parasites for conditional disruption of AMA1 expression

The mCherry sequence was amplified from plasmid pDC2-mCherry-MCS using primers mCherry_PacI_F and mCherry_Bsu361_R. The resulting fragment was digested with Pac I and Bsu361 and cloned into pGEM-T-iA1-C pre-digested with the same enzymes, giving rise to plasmid pGEM-T-ΔA1. PCR screening relied on use of the forward primer A1_Int_F1 in combination with reverse primers A1_EndCont_R1 for the endogenous locus (1.3 kb), A1_Int_R for the integrated locus (1.4 kb) and A1_dsBsu361_R for both non-excised (3.8 kb) and excised (2.0 kb) loci.

#### Guide/Cas9 plasmids

Guide RNA sequences specific for each target gene of interest were identified using Protospacer (www.protospacer.com) or Benchling (www.benchling.com). Selected guide RNA sequences were cloned into plasmid pDC2-Cas9-hDHFRyFCU, containing a Cas9 expression cassette and the drug selection marker human dihydrofolate reductase (*hdhfr*) as previously described ^79^. Complementary oligonucleotides were designed that upon annealing generated sticky ends, compatible with the ends resulting from Bbs1 digestion of plasmid pDC2-Cas9-hDHFRyFCU. To generate conditional-uncleavable forms of AMA1, the *ama1* gene was targeted with guides A1_G1 (oligonucleotides A1-G1_F and A1_G1_R) and A1_g3 (oligonucleotides A1_G3_F and A1_G3_R) which gave rise to plasmids pDC_A1G1 and pDC_A1G3, respectively when cloned into pDC2-Cas9-hDHFRyFCU. Targeting of the *msp1* gene to generate conditional-uncleavable MSP1 was carried out using the *msp1* guide (oligonucleotides msp1guide_F and msp1guide_R) resulting in plasmid pDC_M1G. To generate a conditional knock-out AMA1 line, AMA1guide2 (oligonucleotides AMA1guide2_ and AMA1guide2_R) were cloned into pDC2-Cas9-hDHFRyFCU giving rise to plasmid pDC_A1G2. The U6 cassette from plasmid pDC_A1G2 was amplified using Q5 high fidelity DNA Polymerase (New England BioLabs; NEB) with primers HD104_Cas9UTR_F and HD104_Cas9UTR_R. The resulting PCR product was cloned into pDC_A1G3 pre-digested with Sal I using Gibson Assembly master mix (NEB) according to the manufacturer’s instructions giving rise to plasmid pDC_A1G2/3.

### Oligonucleotide primers used in this study

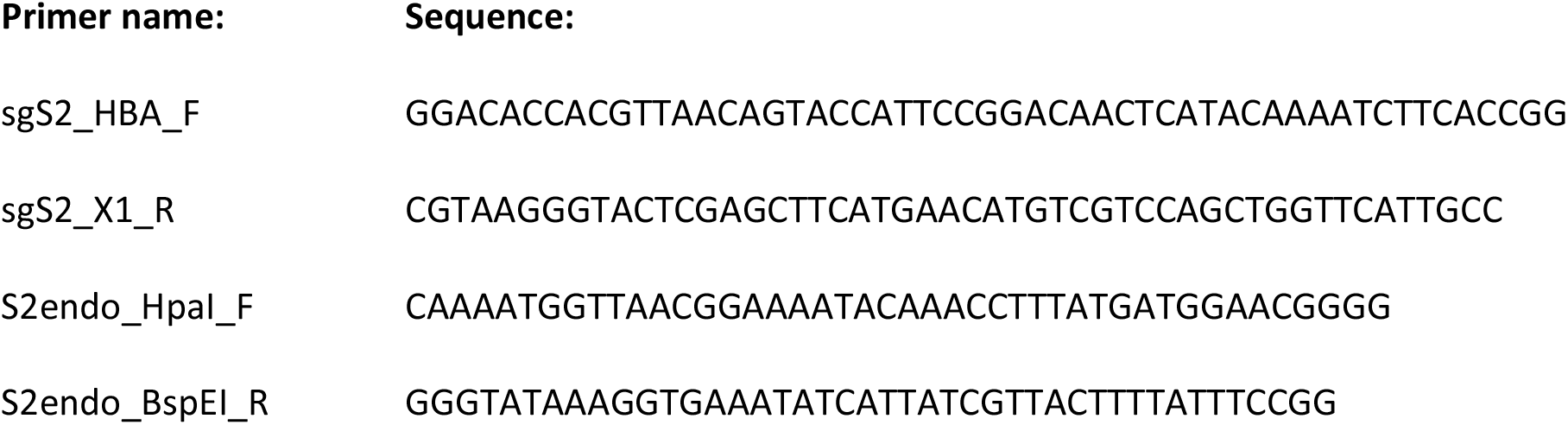
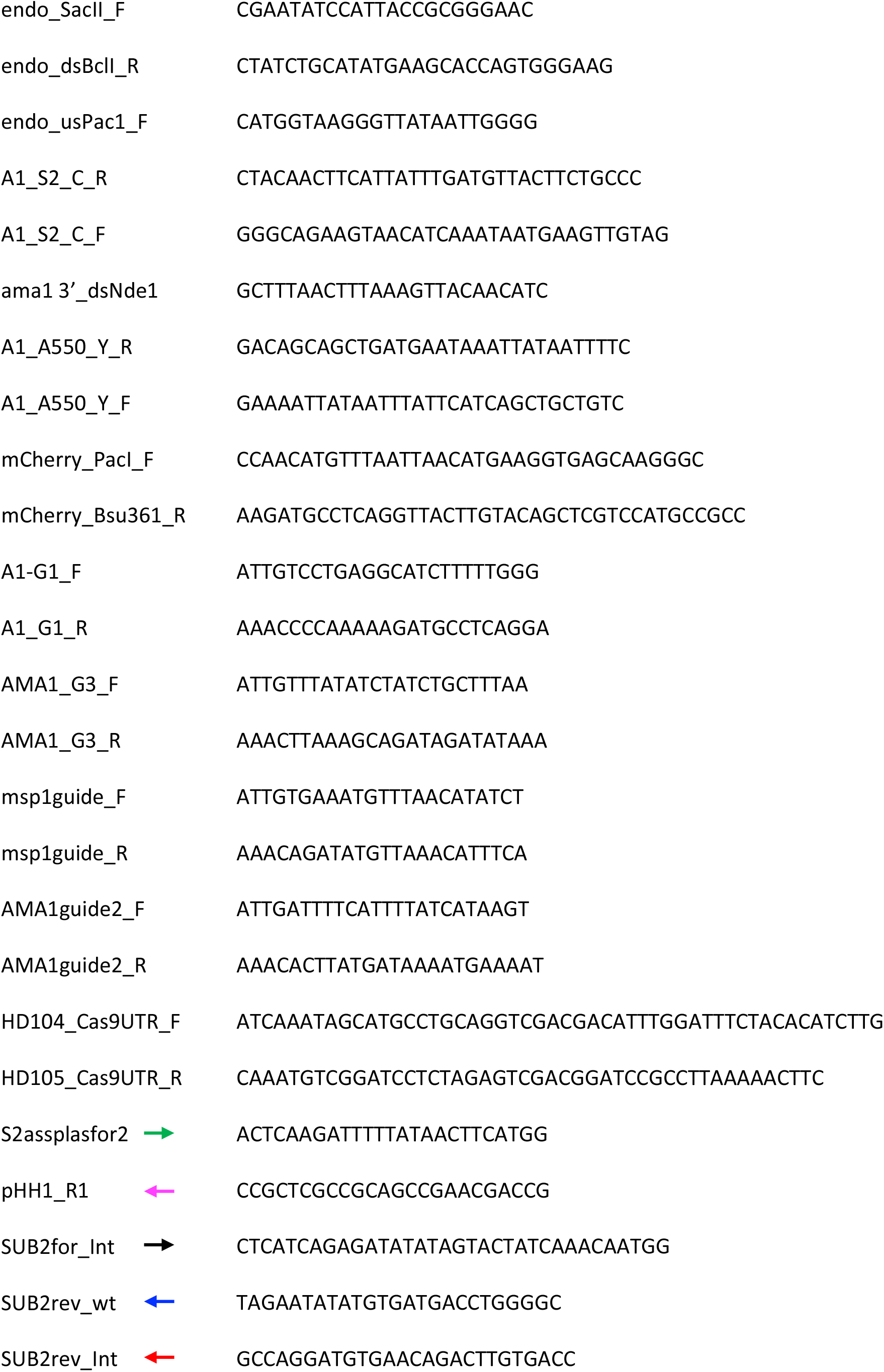
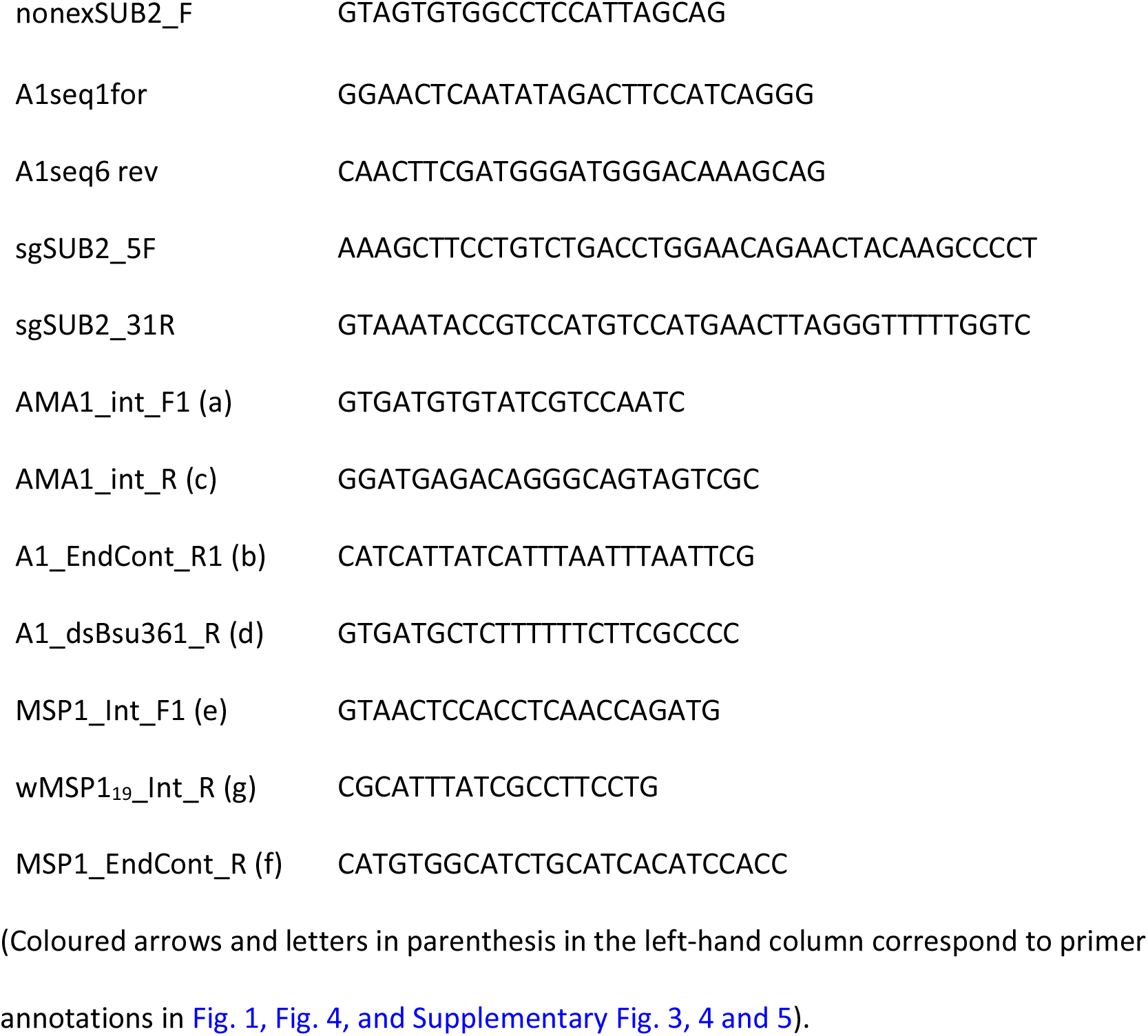

### Generation of transgenic *P. falciparum* lines

Transgenic parasite lines were generated on the background of either the 1G5DC ^28^ or B11 DiCre-expressing parasite lines. In all cases DNA was introduced by electroporation of purified schizonts with sterile DNA, in Amaxa Primary cell solution P3, using a 4D-Nucleofector (Lonza). For single cross-over homologous recombination, 80 μg of purified DNA was used. Drug pressure (WR99210, 2.5 nM; Sigma-Aldrich) was applied 24 h post transfection and maintained until viable parasites were obtained. For stable transgenic lines, parasites were subjected to multiple rounds of drug pressure as previously described ^14^. For Cas9-targeted double cross-over homologous recombination, 20 μg of the guide/Cas9 plasmid and 60 μg of *Sap* I or *Nar* I linearised repair plasmid were used ^79^. Drug selection pressure (2.5 nM WR99210) was applied 24 h following transfection and maintained for 4 d before removal and continued culture of the parasites.

### Plaque assay and limiting dilution cloning of transgenic parasite lines

Plaque assays of transgenic parasites were as described previously ^36^. For limiting dilution cloning, 200μl of culture containing parasites at an estimated 1 parasite per 200 μl and 0.75% haematocrit, was added to each well of a flat-bottomed 96-well plate. Wells containing single plaques were identified after 10-14 days using an inverted microscope and the parasites expanded for further analysis.

### PCR validation of transgenic parasite lines

Parasite pellets were lysed with 0.15% saponin and genomic DNA extracted using a Qiagen DNeasy Blood and Tissue Kit (Qiagen). PCR screens were carried out using GoTaq Green (Promega) with the primer combinations described.

### Invasion and growth assays

For estimation of invasion efficiency, Percoll-enriched schizonts were added to fresh RBCs to obtain a parasitaemia of 5-10% in 4 ml at a haematocrit of 2% haematocrit in RPMI-Albumax. Cultures were incubated at 37°C in a shaking incubator for 4 h. Ring stage parasites were purified as described above and returned to culture in RPMI-Albumax. Samples were taken, fixed with 0.2% glutaraldehyde and stored at 4°C for flow cytometry analysis. The remaining culture was followed for 48 h to check parasite development. Giemsa-stained thin films were prepared as required for microscopic analysis.

For longer term replication assays, cultures were synchronised as described and resulting ring-stage cultures maintained for 24 h to mature to trophozoite stage. Parasitaemia levels were calculated and cultures adjusted to 0.1% parasitaemia, 2% haematocrit in a final volume of 2 ml per well of a 6 well plate. Samples were then taken at t = 0, 48, 96, and 144 h, fixed in 0.2% glutaraldehyde and stored at 4°C for flow cytometry analysis. Culture media were replaced 96 h and 120 h.

### Flow cytometry for parasite quantification

Parasite samples were fixed in 0.2% glutaraldehyde in PBS and stored at 4°C. Cells were prepared for analysis by staining with 2 x SYBR Green I nucleic acid gel stain (Life Technologies) for 30 min at 37°C. Labeling was stopped with an equal volume of PBS and samples analysed using a BD Fortessa flow cytometer (BD Biosciences) with BD FACSDiva software (BD Biosciences). Total RBC numbers were calculated using forward- and side-scatter while fluorescence was detected using the 530/30 blue detection laser. Fluorescence intensity was used to distinguish uninfected from infected RBCs, low fluorescence indicating uninfected cells and gating fixed accordingly. Data were analysed using FlowJo.

### Immunofluorescence analysis (IFA)

Thin films of parasite cultures were made on glass slides, then air dried and stored under dessicant at-80°C. Slides were thawed, fixed in 4% paraformaldehyde for 30 min then permeabilized in 0.1% Triton X-100 in phosphate buffered saline (PBS) for 10 min prior to blocking in 3% (w/v) BSA in PBS overnight. Antibody incubations were carried out for 30 min at 37°C in a humidified chamber followed by washing twice for 5 min each in PBS. All antibodies were diluted into 3% (w/v) BSA in PBS. For microscopic imaging, samples were mounted in Vectashield containing DAPI (Vector laboratories). Images were acquired using a Nikon Eclipse Ni microscope with a 100x Plan Apo λ HA 1.45 objective, with a Hamamatsu C11440 digital camera and processed using Fiji.

### Visualisation of RBC ghost formation using fluorescent phalloidin

Synchronous ring-stage 1B4int parasites were mock or RAP treated, then ∼44 h later schizonts were Percoll-enriched and added to fresh RBCs for 4 h to allow egress and invasion. Without an intervening centrifugation step, a sample of the culture was supplemented with Hoechst 33342 (2 μg/mL; ThermoFisher) and Alexafluor 488 Phalloidin (diluted 1:50; ThermoFisher) and immediately applied to a viewing chamber for live imaging as described previously ^80^. Z-stack images were acquired at 2 μm intervals using a Nikon Eclipse Ni microscope with a 100x Plan Apo λ HA 1.45 objective and a Hamamatsu C11440 digital camera. The number of phalloidin labelled ghosts per image was counted manually for each treatment.

### Haemoglobin release assay

DMSO- or RAP-treated mature 1B4int schizonts were Percoll-enriched and resuspended in fresh RPMI 1640-Albumax. Schizont suspensions were mixed with an equal volume of either RPMI1640-Albumax (control; schizonts only) or RBCs at a 2% haematocrit in RPMI1640-Albumax (co-culture; schizonts plus RBCs). Two additional control samples were set up comprising a similar volume of only RBCs (2% haematocrit) in RPMI1640-Albumax mixed with an equal volume of RPMI1640-Albumax (RBC controls). RBCs from one of these samples were immediately pelleted by centrifugation, the supernatant removed and 200 μl water added to obtain hypotonic lysis of the RBCs. Complete lysis was ensured by subjecting the sample to repeated freeze thawing. RPMI1640-Albumax was then added to return the sample to the original volume. This sample acted as total RBC lysate control.

The schizont only control samples, the co-culture samples and the second RBC control sample were incubated with gentle shaking for 4-6 h at 37°C to allow egress and/or invasion. In some experiments the medium was supplemented with cytochalasin D (10 μM) or 0.5% DMSO (vehicle only) as a control. All samples were then clarified by centrifugation and the culture medium filtered through a 2 μm filter. The samples of culture medium were then subjected to serial 2-fold dilutions in RPMI1640-Albumax, and the haemoglobin content relative to that of the total RBC lysate determined by absorbance at 405 nm ^81^ using a Spectromax M multi-mode microplate reader (Molecular Devices). The pelleted cells from the schizont-containing samples were centrifuged over Percoll cushions to remove residual schizonts and the newly-invaded rings returned to culture. Samples of these were taken immediately as well as 18 h and 24 h later for flow cytometry and microscopic analysis.

### Time-lapse DIC video microscopy of egress and invasion

Visualisation of egress was as described previously ^75^. Briefly, mature schizonts were Percoll-enriched then returned to culture for 4 h in RPMI1640-Albumax supplemented with compound 2 (4-[7-[(dimethylamino)methyl]-2-(4-fluorphenyl)imidazo[1,2-*α*]pyridine-3-yl]pyrimidin-2-amine, C2; 1 μM) to arrest and synchronise the schizonts in a highly mature, pre-egress stage. The parasites were then pelleted, washed twice in pre-warmed, gassed RPMI1640-Albumax medium without C2, then finally pelleted at 1,800 x g for 1 min and immediately resuspended at ∼1 × 10^7^cells/μl in pre-warmed gassed medium without C2. The parasites were introduced by capillary flow into a pre-warmed viewing chamber, made by adhering a 22 × 64 mm borosilicate glass coverslip to a microscope slide ^80^, then at once imaged on a 37°C heated stage on a Zeiss Axio Imager M1 microscope, using a EC Plan-Neofluar 1006/1.3 oil immersion DIC objective fitted with an AxioCam MRm camera. Images were collected at 5 s intervals for 30 mins, annotated and exported as QuickTime movies using Axiovision 3.1.

To image invasion, Percoll-enriched, highly synchronous mature schizonts from RAP- or DMSO-treated cultures were added to fresh RBCs at a 2% haematocrit in pre-warmed, gassed RPMI1640-Albumax to achieve a parasitaemia of ∼10%. The cells were introduced into a pre-warmed viewing chamber, transferred to the heated stage (maintained at 37°C) of a Nikon Eclipse Ni-E wide field microscope and imaged with a Hamamatsu C11440 digital camera using a Nikon N Plan Apo 100x 1.45NA oil immersion objective. DIC images were acquired at 1 s intervals for 30 min, exported as ND2 files and annotated using Fiji (Image J version 2).

### Proteomic analysis

Supernatants of DMSO-and RAP-treated 1B4int schizonts samples following egress and invasion in Albumax-free RPMI 1640 medium were subjected to SDS-PAGE on 16.5% Mini-PROTEIN® Tris-Tricine gels (BioRad). Reduced and alkylated proteins from band slices excised from the entire separation profile were in-gel digested at 37°C overnight with 100 ng trypsin (modified sequencing grade, Promega). Supernatants were dried in a vacuum centrifuge and resuspended in 0.1% TFA. Peptides were loaded onto an Ultimate 3000 nanoRSLC HPLC (Thermo Scientific) coupled to a 2 mm x 0.3 mm Acclaim Pepmap C18 trap column (Thermo Scientific) at 15 μl/min prior to elution at 0.25 μl/min through a 50 cm x 75 μm EasySpray C18 column into an Orbitrap Fusion Lumos Tribrid Mass Spectrometer (Thermo Scientific). A gradient of 2% - 32% acetonitrile in 0.1% formic acid over 45 min was used, prior to washing (80% acetonitrile, 0.1% formic acid) and re-equilibration. The Orbitrap was operated in data-dependent acquisition mode with a survey scan at a resolution of 120,000 from m/z 300-1500, followed by MS/MS in TopS mode. Dynamic exclusion was used with a time window of 20 s. The Orbitrap charge capacity was set to a maximum of 1 × e^6^ ions in 10 ms, whilst the LTQ was set to 1 × e^4^ ions in 100 ms. Raw files were processed using Maxquant 1.3.0.5 and Perseus 1.4.0.11. A decoy database of reversed sequences was used to filter false positives, at a peptide false detection rate of 1%.

### Serial block face scanning EM (SBF-SEM) and transmission EM (TEM)

Mature schizonts enriched from DMSO- and RAP-treated 1B4int cultures were incubated with fresh RBCs for 4 h to allow invasion, then newly-invaded rings isolated as described. Either immediately or following further culture for 20 h, the parasites were washed, fixed with 2.5% glutaraldehyde 4% formaldehyde in 0.1 M phosphate buffer (PB), then further washed in 0.2 M phosphate buffer (PB), embedded in 4% low gelling temperature agarose and cut into 1 mm^3^ blocks ^82^.

For TEM, sample blocks were washed 4 × 15 min in PB and post-fixed in 1% osmium tetroxide/1.5% potassium ferrocyanide for 1 h at 4°C. After further washes in PB at room temperature, the blocks were stained in 1% tannic acid for 45 min and quenched in 1% sodium sulphate for 5 min. The blocks were then washed 4 × 5 min in distilled water and dehydrated through an ethanol series (70-100%, 2 × 10 min each). Finally, the sample blocks were embedded by incubating in propylene oxide (PO) for 10 min, then overnight in a 1:1 PO:Epon 812 (TAAB) resin mixture, followed by 3 changes of pure Epon resin over 2 d, and baking at 60°C for 24 h. For TEM image collection, 80 nm sections were stained with lead citrate and imaged in a Tecnai Spirit BioTwin (Thermofisher Scientific) transmission electron microscope.

For SBF-SEM, samples were embedded in agarose as above, then processed using a protocol adapted from the NCMIR protocol (https://ncmir.ucsd.edu/sbem-protocol). Briefly, the sample blocks were washed 4 × 15 min in 0.1 M PB and post-fixed in 2% osmium tetroxide/1.5% potassium ferrocyanide for 1 h at 4°C, washed 5 × 3 min in water (also after the following steps up to the dehydration), incubated in 1% w/v thiocarbohydrazide for 20 min before a second staining with 2% osmium tetroxide for 30 min, followed by incubation overnight in 1% aqueous uranyl acetate at 4°C. The sample blocks were then stained with Walton’s lead aspartate for 30 min at 60°C and dehydrated through an ethanol series on ice (25-100%, 10 min each). Finally, the sample blocks were embedded as above, except Durcupan (TAAB) resin was used and the baking step was for 48 h. Embedded samples were trimmed and mounted on pins using conductive epoxy glue ^83^. SBF-SEM images were collected using a 3View 2XP system (Gatan Inc), mounted on a Sigma VP scanning electron microscope (Zeiss). Images were collected at 1.9 kV, with a 30 μm aperture, at 6 Pa chamber pressure, and a 2 μs dwell time. The datasets were 69.63 × 69.63 × 13 μm, and 69.63 × 69.63 × 12.6 μm in xyz, consisting of 260 and 252 serial images of 50 nm thickness and a pixel size of 8.5 × 8.5 nm. The total volume of the datasets was ∼63,028 μm^3^ and ∼61,089 μm^3^. SBF-SEM movies were processed in Fiji to adjust brightness and contrast, binned to 4,096 × 4,096 pixels, and then converted to AVI, before exporting from Quicktime Pro as 950 × 950 pixel H.264 movies.

### Quantification and statistical analysis

All graphs of experimental data and statistical analysis were generated using GraphPad Prism 7.0. Statistical analysis methods for each experiment are stated in the corresponding figure legend.

## Lead contact and materials availability

Further information and request for resources and reagents should be directed to and will be fulfilled by corresponding contact Michael J. Blackman. All new plasmids and parasites lines generated in this study can be requested from the corresponding author and will be provided subject to a completed Material Transfer Agreement.

## Ethical statement

Human RBCs were supplied by the UK NHS Blood and Transplant (NHSBT) and approved for use by their internal ethical review process.

## Data availability

All data generated or analysed during this study are included in this published article (and its supplementary information files).

## REFERENCES

1. Aikawa M, Miller LH, Johnson J, Rabbege J. Erythrocyte entry by malarial parasites. A moving junction between erythrocyte and parasite. The Journal of cell biology 77, 72–82 (1978).

2. Dvorak JA, Miller LH, Whitehouse WC, Shiroishi T. Invasion of erythrocytes by malaria merozoites. Science 187, 748–750 (1975).

3. Weiss GE, et al. Revealing the sequence and resulting cellular morphology of receptor-ligand interactions during Plasmodium falciparum invasion of erythrocytes. PLoS Pathog 11, e1004670 (2015).

4. Bannister LH, Butcher GA, Dennis ED, Mitchell GH. Structure and invasive behaviour of Plasmodium knowlesi merozoites in vitro. Parasitology 71, 483–491 (1975).

5. Holder AA, et al. A malaria merozoite surface protein (MSP1)-structure, processing and function. Mem Inst Oswaldo Cruz 87 Suppl 3, 37–42 (1992).

6. Koussis K, et al. A multifunctional serine protease primes the malaria parasite for red blood cell invasion. EMBO J 28, 725–735 (2009).

7. Yeoh S, et al. Subcellular discharge of a serine protease mediates release of invasive malaria parasites from host erythrocytes. Cell 131, 1072–1083 (2007).

8. McBride JS, Heidrich HG. Fragments of the polymorphic Mr 185,000 glycoprotein from the surface of isolated Plasmodium falciparum merozoites form an antigenic complex. Molecular and biochemical parasitology 23, 71–84 (1987).

9. Das S, et al. Processing of Plasmodium falciparum Merozoite Surface Protein MSP1 Activates a Spectrin-Binding Function Enabling Parasite Egress from RBCs. Cell Host Microbe 18, 433–444 (2015).

10. Riglar DT, et al. Super-resolution dissection of coordinated events during malaria parasite invasion of the human erythrocyte. Cell Host Microbe 9, 9–20 (2011).

11. Blackman MJ, Dennis ED, Hirst EM, Kocken CH, Scott-Finnigan TJ, Thomas AW. Plasmodium knowlesi: secondary processing of the malaria merozoite surface protein-1. Exp Parasitol 83, 229–239 (1996).

12. Blackman MJ, Heidrich HG, Donachie S, McBride JS, Holder AA. A single fragment of a malaria merozoite surface protein remains on the parasite during red cell invasion and is the target of invasion-inhibiting antibodies. J Exp Med 172, 379–382 (1990).

13. Blackman MJ, Whittle H, Holder AA. Processing of the Plasmodium falciparum major merozoite surface protein-1: identification of a 33-kilodalton secondary processing product which is shed prior to erythrocyte invasion. Molecular and biochemical parasitology 49, 35–44 (1991).

14. Harris PK, et al. Molecular identification of a malaria merozoite surface sheddase. PLoS Pathog 1, 241–251 (2005).

15. Hackett F, Sajid M, Withers-Martinez C, Grainger M, Blackman MJ. PfSUB-2: a second subtilisin-like protein in Plasmodium falciparum merozoites. Molecular and biochemical parasitology 103, 183–195 (1999).

16. Barale JC, et al. Plasmodium falciparum subtilisin-like protease 2, a merozoite candidate for the merozoite surface protein 1-42 maturase. Proc Natl Acad Sci U S A 96, 6445–6450 (1999).

17. Howell SA, Well I, Fleck SL, Kettleborough C, Collins CR, Blackman MJ. A single malaria merozoite serine protease mediates shedding of multiple surface proteins by juxtamembrane cleavage. J Biol Chem 278, 23890–23898 (2003).

18. Green JL, Hinds L, Grainger M, Knuepfer E, Holder AA. Plasmodium thrombospondin related apical merozoite protein (PTRAMP) is shed from the surface of merozoites by PfSUB2 upon invasion of erythrocytes. Molecular and biochemical parasitology 150, 114–117 (2006).

19. Siddiqui FA, et al. A thrombospondin structural repeat containing rhoptry protein from Plasmodium falciparum mediates erythrocyte invasion. Cell Microbiol 15, 1341–1356 (2013).

20. Thompson J, Cooke RE, Moore S, Anderson LF, Janse CJ, Waters AP. PTRAMP; a conserved Plasmodium thrombospondin-related apical merozoite protein. Molecular and biochemical parasitology 134, 225–232 (2004).

21. Harvey KL, Yap A, Gilson PR, Cowman AF, Crabb BS. Insights and controversies into the role of the key apicomplexan invasion ligand, Apical Membrane Antigen 1. International journal for parasitology 44, 853–857 (2014).

22. Uzureau P, Barale JC, Janse CJ, Waters AP, Breton CB. Gene targeting demonstrates that the Plasmodium berghei subtilisin PbSUB2 is essential for red cell invasion and reveals spontaneous genetic recombination events. Cell Microbiol 6, 65–78 (2004).

23. Zhang M, et al. Uncovering the essential genes of the human malaria parasite Plasmodium falciparum by saturation mutagenesis. Science 360, (2018).

24. Bushell E, et al. Functional Profiling of a Plasmodium Genome Reveals an Abundance of Essential Genes. Cell 170, 260–272 e268 (2017).

25. Olivieri A, et al. Juxtamembrane shedding of Plasmodium falciparum AMA1 is sequence independent and essential, and helps evade invasion-inhibitory antibodies. PLoS Pathog 7, e1002448 (2011).

26. Guevara Patino JA, Holder AA, McBride JS, Blackman MJ. Antibodies that inhibit malaria merozoite surface protein-1 processing and erythrocyte invasion are blocked by naturally acquired human antibodies. J Exp Med 186, 1689–1699 (1997).

27. Lazarou M, et al. Inhibition of erythrocyte invasion and Plasmodium falciparum merozoite surface protein 1 processing by human immunoglobulin G1 (IgG1) and IgG3 antibodies. Infect Immun 77, 5659–5667 (2009).

28. Collins CR, et al. Robust inducible Cre recombinase activity in the human malaria parasite Plasmodium falciparum enables efficient gene deletion within a single asexual erythrocytic growth cycle. Mol Microbiol 88, 687–701 (2013).

29. Howell SA, et al. Distinct mechanisms govern proteolytic shedding of a key invasion protein in apicomplexan pathogens. Mol Microbiol 57, 1342–1356 (2005).

30. Gilson PR, et al. Identification and stoichiometry of glycosylphosphatidylinositol-anchored membrane proteins of the human malaria parasite Plasmodium falciparum. Mol Cell Proteomics 5, 1286–1299 (2006).

31. Ranjan R, et al. Proteome analysis reveals a large merozoite surface protein-1 associated complex on the Plasmodium falciparum merozoite surface. J Proteome Res 10, 680–691 (2011).

32. Lin CS, et al. Multiple Plasmodium falciparum Merozoite Surface Protein 1 Complexes Mediate Merozoite Binding to Human Erythrocytes. J Biol Chem 291, 7703–7715 (2016).

33. Stafford WH, Blackman MJ, Harris A, Shai S, Grainger M, Holder AA. N-terminal amino acid sequence of the Plasmodium falciparum merozoite surface protein-1 polypeptides. Molecular and biochemical parasitology 66, 157–160 (1994).

34. Trucco C, et al. The merozoite surface protein 6 gene codes for a 36 kDa protein associated with the Plasmodium falciparum merozoite surface protein-1 complex. Molecular and biochemical parasitology 112, 91–101 (2001).

35. Pachebat JA, et al. The 22 kDa component of the protein complex on the surface of Plasmodium falciparum merozoites is derived from a larger precursor, merozoite surface protein 7. Molecular and biochemical parasitology 117, 83–89 (2001).

36. Thomas JA, et al. Development and Application of a Simple Plaque Assay for the Human Malaria Parasite Plasmodium falciparum. PloS one 11, e0157873 (2016).

37. Child MA, Harris PK, Collins CR, Withers-Martinez C, Yeoh S, Blackman MJ. Molecular determinants for subcellular trafficking of the malarial sheddase PfSUB2. Traffic 14, 1053–1064 (2013).

38. O’Donnell RA, Preiser PR, Williamson DH, Moore PW, Cowman AF, Crabb BS. An alteration in concatameric structure is associated with efficient segregation of plasmids in transfected Plasmodium falciparum parasites. Nucleic acids research 29, 716–724 (2001).

39. O’Donnell RA, et al. A genetic screen for improved plasmid segregation reveals a role for Rep20 in the interaction of Plasmodium falciparum chromosomes. EMBO J 21, 1231–1239 (2002).

40. van Dijk MR, Vinkenoog R, Ramesar J, Vervenne RA, Waters AP, Janse CJ. Replication, expression and segregation of plasmid-borne DNA in genetically transformed malaria parasites. Molecular and biochemical parasitology 86, 155–162 (1997).

41. Miller LH, Aikawa M, Johnson JG, Shiroishi T. Interaction between cytochalasin B-treated malarial parasites and erythrocytes. Attachment and junction formation. J Exp Med 149, 172–184 (1979).

42. Atkinson MA, Morrow JS, Marchesi VT. The polymeric state of actin in the human erythrocyte cytoskeleton. J Cell Biochem 18, 493–505 (1982).

43. Glushakova S, Humphrey G, Leikina E, Balaban A, Miller J, Zimmerberg J. New stages in the program of malaria parasite egress imaged in normal and sickle erythrocytes. Curr Biol 20, 1117–1121 (2010).

44. Pizarro JC, et al. Crystal structure of the malaria vaccine candidate apical membrane antigen 1. Science 308, 408–411 (2005).

45. Sanders PR, et al. Distinct protein classes including novel merozoite surface antigens in Raft-like membranes of Plasmodium falciparum. J Biol Chem 280, 40169–40176 (2005).

46. Marshall VM, Tieqiao W, Coppel RL. Close linkage of three merozoite surface protein genes on chromosome 2 of Plasmodium falciparum. Molecular and biochemical parasitology 94, 13–25 (1998).

47. Marshall VM, et al. A second merozoite surface protein (MSP-4) of Plasmodium falciparum that contains an epidermal growth factor-like domain. Infect Immun 65, 4460–4467 (1997).

48. Kadekoppala M, Ogun SA, Howell S, Gunaratne RS, Holder AA. Systematic genetic analysis of the Plasmodium falciparum MSP7-like family reveals differences in protein expression, location, and importance in asexual growth of the blood-stage parasite. Eukaryot Cell 9, 1064–1074 (2010).

49. Boyle MJ, et al. Sequential processing of merozoite surface proteins during and after erythrocyte invasion by Plasmodium falciparum. Infect Immun 82, 924–936 (2014).

50. Dluzewski AR, et al. Formation of the food vacuole in Plasmodium falciparum: a potential role for the 19 kDa fragment of merozoite surface protein 1 (MSP1(19)). PloS one 3, e3085 (2008).

51. Volz JC, et al. Essential Role of the PfRh5/PfRipr/CyRPA Complex during Plasmodium falciparum Invasion of Erythrocytes. Cell Host Microbe 20, 60–71 (2016).

52. Suss-Toby E, Zimmerberg J, Ward GE. Toxoplasma invasion: the parasitophorous vacuole is formed from host cell plasma membrane and pinches off via a fission pore. Proc Natl Acad Sci U S A 93, 8413–8418 (1996).

53. Pavlou G, et al. Toxoplasma Parasite Twisting Motion Mechanically Induces Host Cell Membrane Fission to Complete Invasion within a Protective Vacuole. Cell Host Microbe 24, 81–96 e85 (2018).

54. Yahata K, Treeck M, Culleton R, Gilberger TW, Kaneko O. Time-lapse imaging of red blood cell invasion by the rodent malaria parasite Plasmodium yoelii. PloS one 7, e50780 (2012).

55. Alexander DL, Mital J, Ward GE, Bradley P, Boothroyd JC. Identification of the moving junction complex of Toxoplasma gondii: a collaboration between distinct secretory organelles. PLoS Pathog 1, e17 (2005).

56. Srinivasan P, et al. Binding of Plasmodium merozoite proteins RON2 and AMA1 triggers commitment to invasion. Proc Natl Acad Sci U S A 108, 13275–13280 (2011).

57. Lamarque M, et al. The RON2-AMA1 interaction is a critical step in moving junction-dependent invasion by apicomplexan parasites. PLoS Pathog 7, e1001276 (2011).

58. Tyler JS, Boothroyd JC. The C-terminus of Toxoplasma RON2 provides the crucial link between AMA1 and the host-associated invasion complex. PLoS Pathog 7, e1001282 (2011).

59. Lamarque MH, et al. Plasticity and redundancy among AMA-RON pairs ensure host cell entry of Toxoplasma parasites. Nat Commun 5, 4098 (2014).

60. Collins CR, Withers-Martinez C, Hackett F, Blackman MJ. An inhibitory antibody blocks interactions between components of the malarial invasion machinery. PLoS Pathog 5, e1000273 (2009).

61. Giovannini D, et al. Independent roles of apical membrane antigen 1 and rhoptry neck proteins during host cell invasion by apicomplexa. Cell Host Microbe 10, 591–602 (2011).

62. Yap A, et al. Conditional expression of apical membrane antigen 1 in Plasmodium falciparum shows it is required for erythrocyte invasion by merozoites. Cell Microbiol 16, 642–656 (2014).

63. Mital J, Meissner M, Soldati D, Ward GE. Conditional expression of Toxoplasma gondii apical membrane antigen-1 (TgAMA1) demonstrates that TgAMA1 plays a critical role in host cell invasion. Mol Biol Cell 16, 4341–4349 (2005).

64. Bargieri DY, et al. Apical membrane antigen 1 mediates apicomplexan parasite attachment but is dispensable for host cell invasion. Nat Commun 4, 2552 (2013).

65. Parker ML, et al. Dissecting the interface between apicomplexan parasite and host cell: Insights from a divergent AMA-RON2 pair. Proc Natl Acad Sci U S A 113, 398–403 (2016).

66. McNeil PL, Miyake K, Vogel SS. The endomembrane requirement for cell surface repair. Proc Natl Acad Sci U S A 100, 4592–4597 (2003).

67. Stallmach R, et al. Plasmodium falciparum SERA5 plays a non-enzymatic role in the malarial asexual blood-stage lifecycle. Mol Microbiol 96, 368–387 (2015).

68. Withers-Martinez C, et al. Malarial EBA-175 region VI crystallographic structure reveals a KIX-like binding interface. J Mol Biol 375, 773–781 (2008).

69. Roger N, Dubremetz JF, Delplace P, Fortier B, Tronchin G, Vernes A. Characterization of a 225 kilodalton rhoptry protein of Plasmodium falciparum. Molecular and biochemical parasitology 27, 135–141 (1988).

70. Holder AA, Freeman RR. Biosynthesis and processing of a Plasmodium falciparum schizont antigen recognized by immune serum and a monoclonal antibody. J Exp Med 156, 1528–1538 (1982).

71. Burghaus PA, Holder AA. Expression of the 19-kilodalton carboxy-terminal fragment of the Plasmodium falciparum merozoite surface protein-1 in Escherichia coli as a correctly folded protein. Molecular and biochemical parasitology 64, 165–169 (1994).

72. Holder AA, et al. Primary structure of the precursor to the three major surface antigens of Plasmodium falciparum merozoites. Nature 317, 270–273 (1985).

73. Perrin AJ, Collins CR, Russell MRG, Collinson LM, Baker DA, Blackman MJ. The Actinomyosin Motor Drives Malaria Parasite Red Blood Cell Invasion but Not Egress. mBio 9, (2018).

74. Blackman MJ. Purification of Plasmodium falciparum merozoites for analysis of the processing of merozoite surface protein-1. Methods Cell Biol 45, 213–220 (1994).

75. Thomas JA, et al. A protease cascade regulates release of the human malaria parasite Plasmodium falciparum from host red blood cells. Nat Microbiol 3, 447–455 (2018).

76. Child MA, Epp C, Bujard H, Blackman MJ. Regulated maturation of malaria merozoite surface protein-1 is essential for parasite growth. Mol Microbiol 78, 187–202 (2010).

77. Kocken CH, et al. High-level expression of the malaria blood-stage vaccine candidate Plasmodium falciparum apical membrane antigen 1 and induction of antibodies that inhibit erythrocyte invasion. Infect Immun 70, 4471–4476 (2002).

78. Collins CR, Withers-Martinez C, Bentley GA, Batchelor AH, Thomas AW, Blackman MJ. Fine mapping of an epitope recognized by an invasion-inhibitory monoclonal antibody on the malaria vaccine candidate apical membrane antigen 1. J Biol Chem 282, 7431–7441 (2007).

79. Knuepfer E, Napiorkowska M, van Ooij C, Holder AA. Generating conditional gene knockouts in Plasmodium - a toolkit to produce stable DiCre recombinase-expressing parasite lines using CRISPR/Cas9. Sci Rep 7, 3881 (2017).

80. Collins CR, et al. Malaria parasite cGMP-dependent protein kinase regulates blood stage merozoite secretory organelle discharge and egress. PLoS Pathog 9, e1003344 (2013).

81. Snell SM, Marini MA. A convenient spectroscopic method for the estimation of hemoglobin concentrations in cell-free solutions. J Biochem Biophys Methods 17, 25–33 (1988).

82. Hanssen E, Goldie KN, Tilley L. Ultrastructure of the asexual blood stages of Plasmodium falciparum. Methods Cell Biol 96, 93–116 (2010).

83. Russell MR, et al. 3D correlative light and electron microscopy of cultured cells using serial blockface scanning electron microscopy. J Cell Sci 130, 278–291 (2017).

