## Supplemental Information for "The malaria parasite sheddase SUB2 governs host red blood cell membrane sealing at invasion"

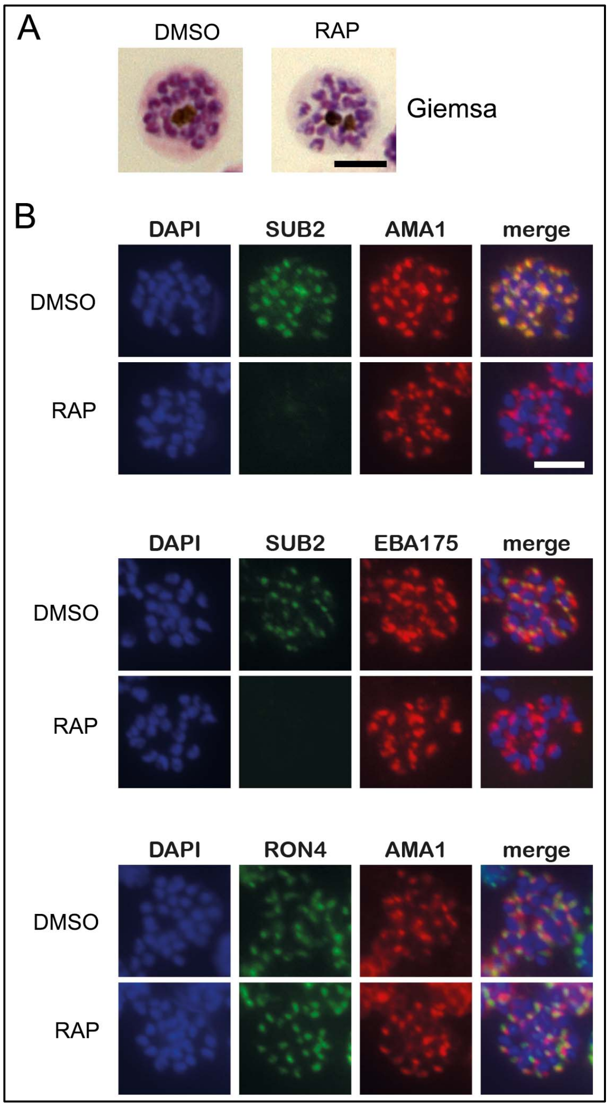


**Supplementary Figure 1. Related to Figure 1. Loss of SUB2 expression in cycle 0 has no impact on schizont development or morphology**

(A) Light micrographs of Giemsa-stained DMSO- and rapamycin (RAP)-treated 1B4int schizonts (cycle 0) showing normal morphology in both cases.

(B) IFA of DMSO- (control) and RAP-treated mature cycle 0 1B4int schizonts, showing the expected punctate localisation of the microneme proteins apical membrane antigen 1 (AMA) and EBA175, and the rhoptry neck protein RON4. SUB2 was detected with antibodies to the HA3 epitope tag. Parasite nuclei were stained with the DNA dye 4,6-diamidino-2-phenylindole (DAPI, blue). Scale bars, 5 µm.

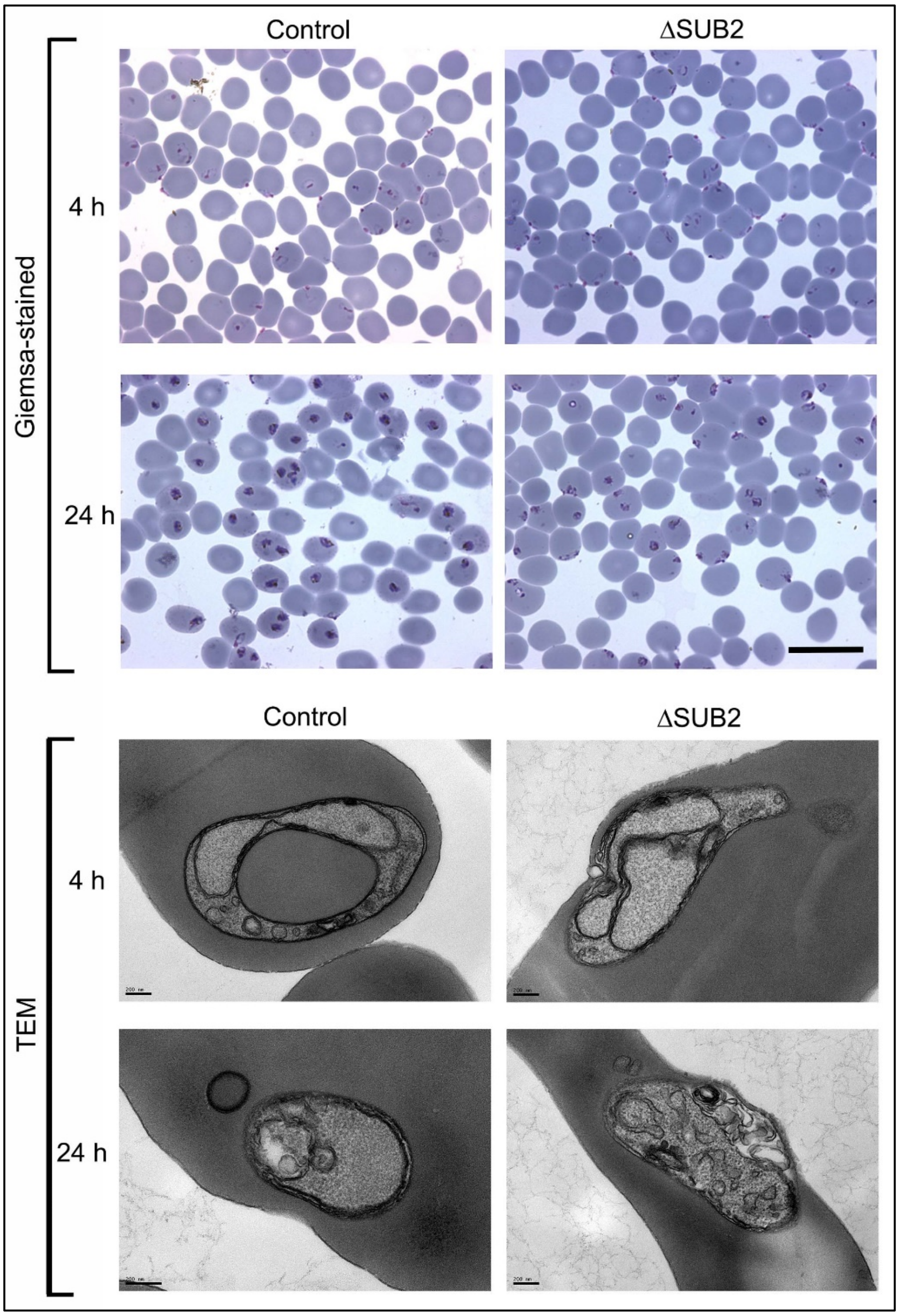


**Supplementary Figure 2. Related to Figure 4. Light and transmission electron microscopic (TEM) analysis of newly-invaded intraerythrocytic ΔSUB2 cycle 1 parasites shows no discernible structural defect**

(A) Light micrographs of Giemsa-stained cycle 1 rings and trophozoites sampled 4 h and 24 h following invasion by control or RAP-treated (ΔSUB2) 1B4int parasites. The 4 h rings are morphologically indistinguishable, but retarded development of the ΔSUB2 parasites is evident by 24 h. Scale bar, 20 μm.

(B) Electron micrographs of parasite-infected cells taken at similar time points. Note that whilst the 4 h ΔSUB2 ring appears to display a potential sealing defect, similar structures were observed with similar frequency in control ring preparations, and so could not be confidently ascribed as being responsible for the developmental defects in the mutants.

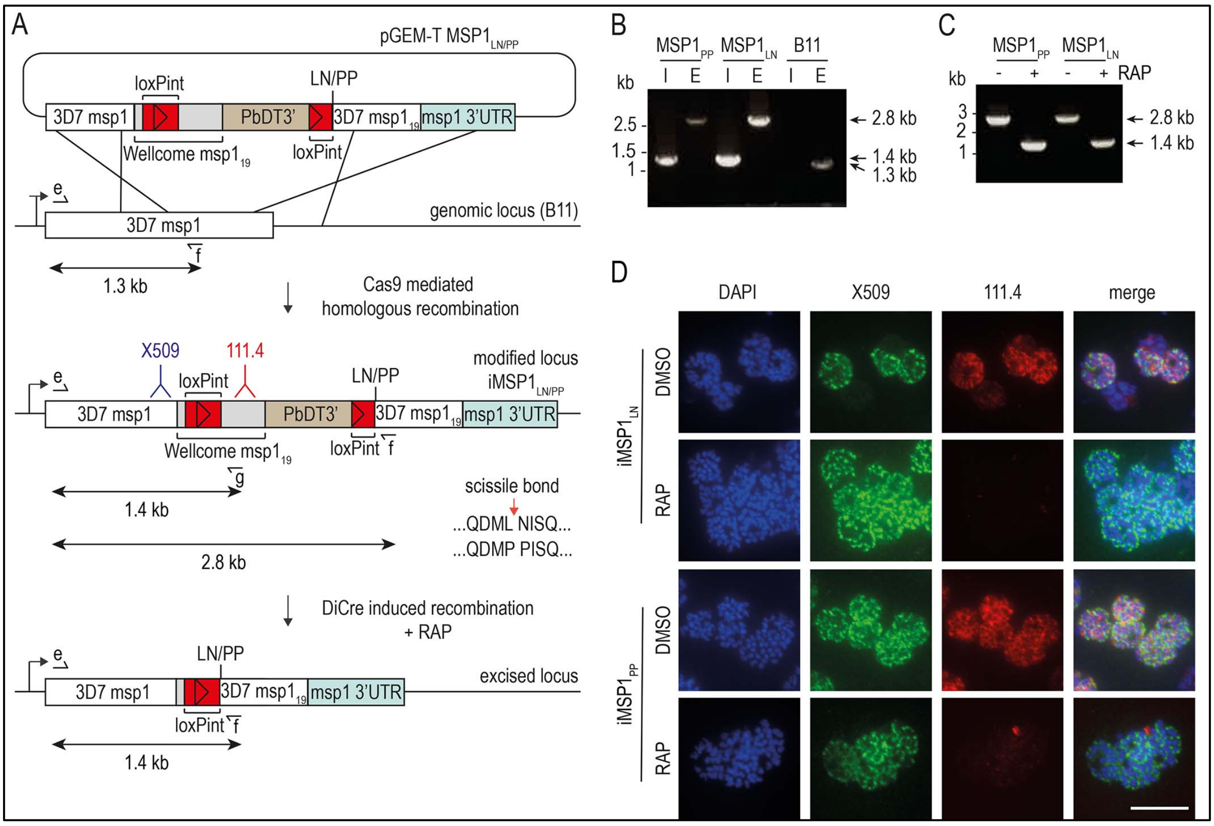


**Supplementary Figure 3. Related to Figure 6. Conditional mutagenesis of the *P. falciparum msp1* gene to selectively block SUB2-mediated shedding of MSP1**

(A) Strategy for modifying the *msp1* locus by Cas9-enhanced homologous recombination in the DiCre-expressing B11 *P. falciparum* clone (Perrin et al., 2018) to generate conditional allelic replacement lines. The targeting construct is depicted above the *msp1* locus both before and following modification, as well as following DiCre-mediated excision of the floxed sequences. Introduced *loxPint* sites, red. Sequence encoding the Wellcome-type $MSP1_{19}$ (grey) and 3D7 $MSP1_{19}$ (white) are indicated (see Methods). Epitopes recognized by monoclonal antibodies (mAbs) X509 (blue) and 111.4 (red) are indicated. Following excision, parasites express modified forms of MSP1 expressing either Leu-Asn or Pro-Pro flanking the SUB2 cleavage site. Primers for diagnostic PCR (see panels B and C) are indicated (half arrows; see Methods for all primer codes and sequences). Predicted sizes of amplicons are shown.

(B) Diagnostic PCR confirmation of gene modification in clonal lines. PCR with primers e plus f, designed to detect the unmodified endogenous locus, produced a 1.3 kb amplicon in the B11 parental line (E) but a 2.8 kb amplicon in the $iMSP1_{LN}$ and $iMSP1_{PP}$ lines. Integration-specific diagnostic PCR (I; primers e plus g) only recognized the modified locus in the $iMSP1_{LN}$ and $iMSP1_{PP}$ lines, generating a 1.4 kb amplicon.

(C) Validation by diagnostic PCR of efficient RAP-induced excision in clonal lines, using primers e plus f, producing an amplicon of 1.4 kb only following excision.

(D) IFA of mature schizonts confirming allelic exchange following RAP-induced recombination in the $iMSP1_{LN}$ and $iMSP1_{PP}$ lines. This results in loss of reactivity with mAb 111.4 which recognises a polymorphic epitope only in the Wellcome-type $MSP1_{19}$ sequence (Child et al., 2010). Scale bar, 10 μm.

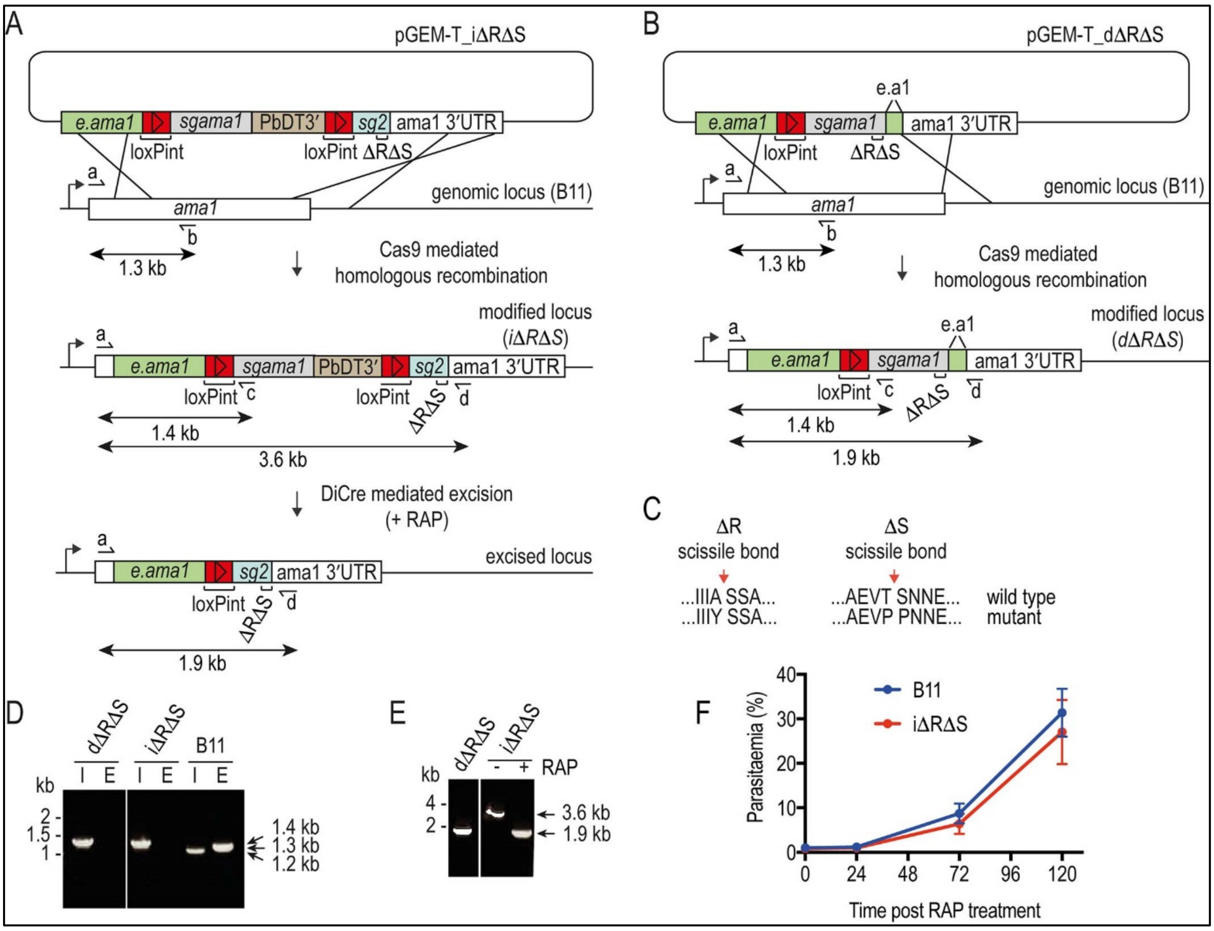


**Supplementary Figure 4. Related to Figure 7. Conditional or direct mutagenesis of the *P. falciparum ama1* gene to selectively block proteolytic shedding of AMA1**

(A) and (B) Schematic of Cas9-enhanced generation of iΔRΔS and dΔRΔS parasite lines in the DiCre-expressing B11 *P. falciparum* line. Recodonised *ama1* gene sequences, grey and blue; endogenous sequence, green; *loxPint* sequences, red. Primers used for diagnostic PCR, half arrows. see Methods for all primer codes and sequences.

(C) Modifications made to the SUB2 and ROM4 cleavage sites to render them refractory to cleavage by the respective proteases.

(D) Diagnostic PCR confirming successful modification of the *ama1* locus in the iΔRΔS and dΔRΔS parasite clones. I, integration-specific PCR (primers a plus c, amplicon size 1.4 kb). E, endogenous (unmodified) locus-specific signal (primers a plus b, amplicon size 1.3 kb). Note that mis-priming in the parental B11 line generates a smaller 1.2 kb product in the integration-specific PCR.

(E) Confirmation of RAP-mediated excision (in the iΔRΔS line) or integration (in the dΔRΔS line) by diagnostic PCR, using primers a plus d. Excision results in reduction of the amplicon from 3.6 kb to 1.9 kb.

(F) Replication rates of the (untreated) iΔRΔS line and B11 parental line are similar.

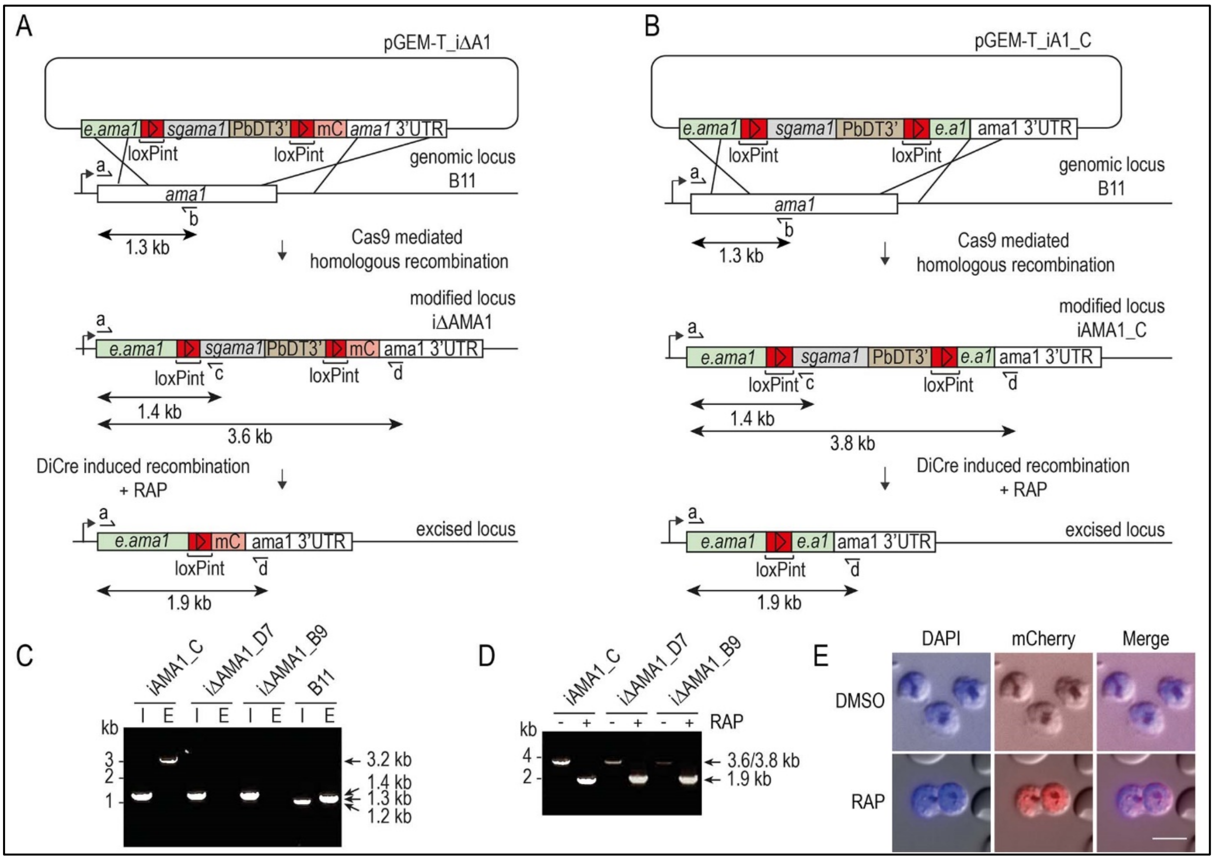


**Supplementary Figure 5. Related to Figure 7. Conditional disruption of the *P. falciparum ama1* gene**
(A) and (B) Schematic representation of Cas9-enhanced generation of inducible disruption iΔAMA1 and control iAMA1_C parasite lines in the DiCre-expressing B11 *P. falciparum* line, and the expected consequences of DiCre-mediated excision. Recodonised *ama1* sequences, grey; endogenous *ama1* sequences, green; *loxPint* sequences, red; mC, mCherry sequence, pink. In (A), DiCre-mediated excision leads to fusion of a truncated AMA1 gene product to mCherry via a reconstituted *loxPint* intron. Primers used for diagnostic PCR, half arrows, with predicted amplicon sizes indicated. see Methods for primer codes and sequences.
(C) Diagnostic PCR confirming successful modification of the *ama1* locus in the control iAMA1_C line and inducible disruption (iΔAMA1_D7 and iΔAMA1_B9) parasite clones. E, endogenous (unmodified) locus-specific signal (primers a plus b, amplicon size 1.3 kb). I, integration-specific PCR (primers a plus c, amplicon size 1.4 kb). Mis-priming in the integration-specific PCR from the parental B11 line generates a smaller 1.2 kb product. The 3.2 kb band in the iAMA1_C endogenous track is due to primer b also hybridising to e.a1, the part of the endogenous *ama1* sequence used as the 3' target in pGEM-T_iA1_C.
D) Diagnostic PCR confirming efficient RAP-mediated excision in all clonal lines. Primers a plus d were used to screen for excision, producing a 3.8 kb or 3.6 kb amplicon from the non-excised loci which reduced to 1.9 kb in both cases upon excision.
(E) Expression of mCherry in schizonts upon RAP-treatment of iΔAMA1_D7 parasites. Scale bar, 10 μm.

**Supplementary Table 1. Related to Figure 3. Quantitative mass spectrometric analysis of *P. falciparum* proteins in egress supernatants of ΔSUB2 1B4int schizonts compared with control schizonts**
Data shown are from 3 technical replicates.

**Supplementary Table 2.** Reagents, cells lines and software used in this study.

**Supplementary Video 1. Related to Figure 4. Serial block face scanning electron microscopy (SBF-SEM) of newly-invaded rings (control)**
Composite video showing SBF-SEM analysis of newly-invaded (4h-old) rings formed following invasion by merozoites released from DMSO-treated (control) cycle 0 1B4int schizonts.

**Supplementary Video 2. Related to Figure 4. Serial block face scanning electron microscopy (SBF-SEM) of newly-invaded rings (ΔSUB2)**
Composite video showing SBF-SEM analysis of newly-invaded (4h-old) rings formed following invasion by merozoites released from RAP-treated (ΔSUB2) cycle 0 1B4int schizonts.

**Supplementary Video 3. Related to Figure 5. Abortive invasion by ΔSUB2 merozoites leads to rapid target RBC lysis**
Time-lapse video microscopy showing interactions between naturally-released merozoites and target human RBCs. For both control and ΔSUB2 merozoites, characteristic RBC deformation during the initial interaction and transient echinocytosis immediately following merozoite entry is observed. In the case of the ΔSUB2 merozoites, lysis of two targeted RBCs (arrowed) occurs at ~740 seconds, as indicated by sudden loss of differential interference contrast (DIC).

**Supplementary Video 4. Related to Figure 7. Interactions between ΔAMA1 merozoites and target RBCs lead to RBC lysis**
Time-lapse video microscopy showing interactions between target RBCs (arrowed) and merozoites released from RAP-treated iΔAMA1 schizonts. Characteristic RBC deformation during the initial interaction and RBC echinocytosis is observed. The target RBCs remained rounded and eventually lysed (see main manuscript Figure 7H) as indicated by loss of DIC.
